# The commitment of barley microspores into embryogenesis involves miRNA-directed regulation of members of the SPL, GRF and HD-ZIPIII transcription factor families

**DOI:** 10.1101/2020.06.11.146647

**Authors:** Sébastien Bélanger, Patricia Baldrich, Marc-André Lemay, Suzanne Marchand, Patricio Esteves, Blake C. Meyers, François Belzile

## Abstract

Microspore embryogenesis is a model for developmental plasticity and cell fate decisions. To investigate the role of miRNAs in this development, we sequenced sRNAs and the degradome of barley microspores collected prior to (day 0) and after (days 2 and 5) the application of a stress treatment known to induce embryogenesis. Microspores isolated at these timepoints were uniform in both appearance and in their complements of sRNAs. We detected 68 miRNAs in microspores. The abundance of 51 of these miRNAs differed significantly during microspore development. One group of miRNAs was induced when the stress treatment was applied, prior to being repressed when microspores transitioned to embryogenesis. Another group of miRNAs were up-regulated in day-2 microspores and their abundance remained stable or increased in day-5 microspores, a timepoint at which the first clear indications of the transition towards embryogenesis were visible. Collectively, these miRNAs might play a role in the modulation of the stress response, the repression of gametic development, and/or the gain of embryogenic potential. A degradome analysis allowed us to validate the role of miRNAs in regulating 41 specific transcripts. We showed that the transition of microspores toward the embryogenesis pathway involves miRNA-directed regulation of members of the *ARF, SPL, GRF* and *HD-ZIPIII* transcription factor families. We noted that 41.5% of these targets were shared between day-2 and day-5 microspores while 26.8% were unique to day-5 microspores. The former set may act to disrupt transcripts involved in pollen development while the latter set may drive the commitment to embryogenesis.

## INTRODUCTION

Gametic embryogenesis (also called microspore embryogenesis or androgenesis) occurs when the microspore, a haploid and uninucleate precursor to the pollen grain, switches from a gametophytic to embryonic fate. This developmental transition is artificially triggered through an inductive stress treatment (Seifert et al., 2016). Microspore embryogenesis is enabled by the totipotency of plant cells and it is a model for developmental plasticity and cell fate decisions in plants (Soriano et al., 2013; Seifert et al., 2016). In the past decades, gametic embryogenesis has been exploited in research and plant breeding to obtain double haploid (DH) plants, as the initially haploid embryos can spontaneously double their chromosomal complement or be induced to do so (Seifert et al., 2016). DH technology is currently employed in many breeding programs in various crop species because it can quickly generate genetically fixed (homozygous), recombinant plants directly from an F1 hybrid (Germanà, 2011; Seifert et al., 2016). Unfortunately, the DH protocol often needs to be developed or fine-tuned on a case-by-case basis, due to a large variability in responsiveness to embryogenesis from one species to another, and even from one genotype to another within a given species (Soriano et al., 2013). One of the major bottlenecks in the process is the lack, or low efficiency, of the switch to the embryogenic pathway (Soriano et al., 2013).

It is assumed that stresses induce an interruption in transcriptional and translational activities which normally lead to pollen formation. This might then restore cell totipotency (due to cellular de-differentiation), and ultimately drive the transition to embryogenesis (Maraschin et al., 2005; Elhiti et al., 2013; Seifert et al., 2016). In other words, for the process to occur, a competent microspore has to stop the expression of genes involved in pollen development, possibly degrade the existing transcripts, and then induce expression of the genes required for commitment to embryogenesis. Although there have been investigations into epigenetic modifications that modulate transcriptional regulation, such as DNA methylation (El-Tantawy et al., 2014), histone methylation (Berenguer et al., 2017) or acetylation (Li et al., 2014), these processes would not be expected to eliminate existing transcripts. Small regulatory RNAs (sRNAs) can accomplish this role since they can degrade and eliminate target transcripts. Furthermore, sRNAs are known to play important roles in key developmental processes such as patterning of the embryo and meristem, leaf, and flower development (D’Ario et al., 2017). However, the role(s) of sRNAs in gametic embryogenesis remains unclear.

Endogenous sRNAs are 21 to 24-nucleotide (nt) RNAs involved in transcriptional gene silencing (TGS) or post-transcriptional gene silencing (PTGS) (Borges and Martienssen, 2015; Komiya, 2017; Axtell and Meyers, 2018). The 24-nt sRNAs are known to be involved in TGS through the RNA-directed DNA methylation (RdDM) pathway, enabling the establishment and maintenance of DNA methylation (Bond and Baulcombe, 2014). Commonly named “heterochromatic small interfering RNAs” (hc-siRNAs), these sRNAs are known to suppress transposable elements (TEs) and repetitive regions (Matzke et al., 2009; Komiya, 2017). Thus, hc-siRNAs may play an important role in the early stages of development through embryogenesis via reprogramming the genomes of microspores and preserving genome integrity during chromosome doubling or the first nuclear division of microspores.

With a typical length of 21- or 22-nt, microRNAs (miRNAs) function in PTGS by facilitating the degradation or translational inhibition of mRNA molecules with complementary sites (D’Ario et al., 2017; Komiya, 2017). miRNAs have an important role in developmental phase changes since they often direct cleavage of transcripts encoding transcription factors (TFs), thus broadly affecting gene regulatory networks (Jones-Rhoades et al., 2006). Since TFs can be involved in developmental patterning or stem cell identity, miRNAs were proposed to play a role in differentiation by targeting transcripts of regulatory genes responsible for existing expression programs, thereby facilitating more rapid and robust transitions to new expression programs (Rhoades et al., 2002; Jones-Rhoades et al., 2006). Such miRNA-assisted reprogramming could be a powerful mechanism to direct the microspore developmental transition from gametic to embryogenic cell fate during isolated microspore culture.

Most plant miRNAs perform their repressive regulation through target site cleavage; the sliced sites of target transcripts can be identified by sequencing the 3’ remnants of cleavage (Shao et al., 2012). Degradome sequencing, also called parallel analysis of RNA ends (PARE; German et al., 2008; Zhai et al., 2014) or genome-wide mapping of uncapped and cleaved transcripts (GMUCT; Addo-Quaye et al., 2008), allows transcriptome-wide mapping of target cleavage sites by combining a modified 5’-RACE approach with next-generation sequencing technologies (Shao et al., 2012).

Barley (*Hordeum vulgare* ssp. *vulgare*) is considered a model species for molecular analysis of microspore embryogenesis in monocots (Soriano et al., 2013). We recently published an RNA-seq analysis of microspore embryogenesis in barley (Bélanger et al., 2018) on a cultivar (Gobernadora) known to exhibit an exceptionally good response to androgenesis (Marchand et al., 2008). In the work described here, we designed our experiments to address perceived limitations in the previous work, namely by concurrently performing both sRNA sequencing (sRNA-seq) and Parallel Analysis of RNA End sequencing (PARE-seq) for the same set of treatments on which RNA-seq alone was performed previously (Bélanger et al., 2018). This paper provides an extensive resource for functional genomics analyses on the induction of microspore embryogenesis in barley.

## MATERIAL AND METHODS

### Microspore production, cellular fixation and microscopy

Donor barley plants (*H. vulgare* ssp. *vulgare* cv. Gobernadora, a two-row spring barley) were grown in a greenhouse. Uniform immature spikes containing microspores at the mid-late to late-uninucleate stage were harvested as described by Esteves and Belzile (2014). We performed the analyses described below at three time points: day 0 (from freshly harvested spikes), day 2 (immediately after completion of the stress pretreatment) and day 5 (after three days in culture) as detailed in Bélanger et al. (2018). For samples at day 0, microspores were isolated from freshly harvested spikes containing haploid and uninucleate microspores. The uniformity of these microspores was improved by performing gradient centrifugation (20% maltose-mannitol; 900x*g* at 12°C). For samples at days 2 and 5, spikes were first subjected to a 48-h pretreatment combining thermal (26°C), osmotic (0.3M mannitol; pH at 5.34) and starvation stresses. After the pretreatment, microspores were harvested and purified via gradient centrifugation (as above). A fraction of these microspores was collected to serve as day-2 samples (i.e. immediately after completion of the pretreatment) while the rest was plated on a two-layer (solid-liquid) embryogenesis-induction medium (Li and Devaux, 2003; Esteves et al., 2014) and cultivated at 28°C for three days. Finally, to maximize the uniformity of the microspores harvested on day 5, we performed another gradient centrifugation (25% maltose-mannitol; 300x*g*; 12°C). The microspores were produced in four replicates and, after isolation, samples were immediately frozen in liquid nitrogen and kept at −80°C prior to RNA isolation.

Additional samples of microspores were collected, fixed and DAPI stained for microscopic analysis as described by González-Melendi et al. (2005) except that the washing of microspores was performed for 15 minutes twice. Microscopy was performed at the Plate-forme d’Imagerie Moléculaire et de Microscopie of the Institut de Biologie Intégrative et des Systèmes (Université Laval, Québec, Canada) using 10 μl of stained microspores and observed with a Zeiss Axio Observer.Z1 (Zeiss, Gottingen, Germany) under a UV laser (excitation of 390/22 nm and emission of 460/50 nm).

### RNA isolation, library construction and sequencing

Total RNA was isolated using the TriZol Reagent Solution (Applied Biosystems, Foster City, CA, USA) as per the manufacturer’s instructions. RNA quality was evaluated using the Agilent RNA 6000 Nano Kit on the Bioanalyser 2100 (Agilent Technologies, Santa Clara, CA, USA). Only samples with an RNA integrity number ≥ 7.0 were kept for library construction. All samples were quantified using the DS-11 FX+ fluorometer (DeNovix, Wilmington, DE, USA). A total of 12 sRNA libraries were constructed from 5 μg of size-selected RNA using the NEBNext^®^ Small RNA Library Prep kit (New England Biolabs, Ipswich, MA, USA) as per the manufacturers’ instructions. Then, total RNAs were combined to form an equimolar composite of all replicates corresponding to microspores at each stage of development. A total of 20 μg of these RNA composites was used to prepare PARE-seq libraries as described by Zhai et al. (2014). Finally, 50-nt single-end sequencing was performed for these two types of RNA libraries (2 and 0.5 lanes for sRNA and PARE libraries, respectively) on an Illumina HiSeq 2000 at the University of Delaware DNA Sequencing & Genotyping Center (Newark, DE, USA).

### Bioinformatics analysis of sRNA-seq data

Using cutadapt v2.9 (Martin, 2011), sRNA-seq reads were pre-processed to remove adapters (Supplementary Table 1) and discard reads shorter than 15 nt. Clean reads were mapped to the barley reference genome v2 (Mascher et al., 2017; available on Ensembl release-44) using ShortStack v3.8.5 (Johnson et al., 2016) with the following parameters: -mismatches 0, -bowtie_m 50, -mmap u, -dicermin 19, -dicermax 25 and -mincov 0.5 transcripts per million (TPM). Results generated by ShortStack were filtered to keep only clusters having a predominant RNA size observed between 20 and 24 nucleotides, inclusively. We then annotated categories of microRNA (miRNA), phased small interfering RNA (phasiRNA) and heterochromatic small interfering RNA (hc-siRNA).

First, sRNA reads representative of each cluster were aligned to the monocot-related miRNAs listed in miRBase release 22 (Kozomara and Griffiths-Jones, 2014; Kozomara et al., 2019) using NCBI BLASTN v2.9.0+ (Comacho et al., 2009) with the following parameters: -strand both, -task blastn-short, -perc_identity 75, -no_greedy and -ungapped. Homology hits were filtered and sRNA reads were considered as known miRNA based on the following criteria: (i) no more than four mismatches and (ii) no more than 2-nt extension or reduction at the 5’ end or 3’ end. Known miRNAs were summarized by family and when genomes contained multiple miRNA loci per family, miRNAs were ordered per chromosomal position and renamed based on these genomic positions. Small RNA reads with no homology to known miRNAs were annotated as novel miRNAs using the *de novo* miRNA annotation performed by ShortStack. The secondary structure of new miRNA precursor sequences was drawn using the RNAfold v2.1.9 program (Lorenz et al., 2011). Candidate novel miRNAs were manually inspected and only those meeting criteria for plant miRNA annotations (Axtell and Meyers, 2018) were kept for downstream analyses. Then, the remaining sRNA clusters were analyzed to identify phasiRNAs based on ShortStack analysis reports. sRNA clusters having a “Phase Score” ≥30 were considered as true positive phasiRNAs. Genomic regions corresponding to these phasiRNAs were considered as *PHAS* loci and grouped in categories of 21- and 24-*PHAS* loci based on the length of phasiRNAs derived from these loci. Non-phased sRNAs having a length of 24-nt were considered as hc-siRNAs.

Finally, the read-count matrix generated by ShortStack was used to perform an expression analysis for annotated sRNAs (miRNA, phasiRNA and hc-siRNA) and to identify those that were differentially expressed over the course of microspore development. Prior to preforming the expression analysis, the read-count matrix was normalized for library size using the TMM method of the edgeR program (Robinson et al., 2010; Robinson and Oshlack, 2010). To assess the degree of uniformity among replicates of the three developmental stages, we performed an MDS analysis using edgeR. Differentially expressed sRNAs were identified using the generalized linear model (glm) test function of edgeR for developmental transitions: (i) from day 0 to day 2, (ii) from day 2 to day 5 and (iii) from day 0 to day 5. Only sRNAs having both a log_2_FC ≥ |2.0| and a *q-value* ≤ 1.0E-03 were considered as differentially expressed. Differentially expressed sRNAs were visualized by heatmap using the R program pheatmap (https://rdrr.io/cran/pheatmap/).

### Bioinformatics analysis of PARE-seq data

To identify miRNA-target pairs in microspores, we performed a degradome analysis for miRNAs annotated in microspores through the sRNA-seq experiment. Using cutadapt v2.9, PARE-seq reads were pre-processed to remove adapters (Supplementary Table 1) and discard reads shorter than 15 nt. Then, we used PAREsnip2 (Thody et al., 2018) to predict all sRNA-target pairs and to validate the effective sRNA-guided cleavage site with PARE-seq reads. We ran PAREsnip2 with default parameters using Fahlgren & Carrington targeting rules (Fahlgren and Carrington, 2010). We considered only targets in categories 0, 1 and 2 for downstream analysis. The gene annotations available on Phytozome V13 and Ensembl Plant release-44 were used to identify *A. thaliana* and *O. sativa* orthologs. Functional annotation of orthologous genes was used to interpret the role of miRNA in microspores.

### Data availability

The complete set of raw sRNA-seq and PARE-seq reads were deposited in the Sequence Read Archive under SRA accession number PRJNA634514.

## RESULTS AND DISCUSSION

### Microspores collected at three key stages capture the switch to embryogenesis

To study the role of sRNAs during the switch from gametophytic to embryonic development, we collected microspores prior to (day 0) or following the application of a pretreatment intended to induce embryogenesis (days 2 and 5) in the barley cultivar cv. Gobernadora. This genotype is known to be highly responsive to gametic embryogenesis (Marchand et al., 2008). We observed, with and without DAPI staining, that (i) day-0 microspores were characterized by having a single nucleus positioned close to the cell wall (Figure 1a); this corresponds to the late uninucleate stage known as the most embryogenic-responsive stage in barley (Kasha et al., 2001a); (ii) day-2 microspores primarily showed a single nucleus migrating toward the center of the cells while a few had two nuclei (Figure 1b), and (iii) day-5 microspores exhibited multiple nuclei (Figure 1c). The microspore populations were homogenous and few or no damaged or dead cells in all stages of development. The phenotypes of microspores on days 2 and 5 were very similar to those described in previous work in barley (Kasha et al., 2001b; Maraschin, 2005). Using a similar protocol, Maraschin (2005) described that a microspore was capable of releasing an embryo-like structure out of the microspore exine wall when microspores initiated nuclear divisions. Since the phenotype of most microspores indicates that nuclear divisions were initiated on day 5, we consider that, by that time, microspores have embarked on a sporophytic development leading to embryogenesis. Our observed phenotypes were highly similar (practically indistinguishable) from those described by Bélanger et al. (2018), thus suggesting an experimental reproducibility in the preparation of these materials.

**Figure 1.**
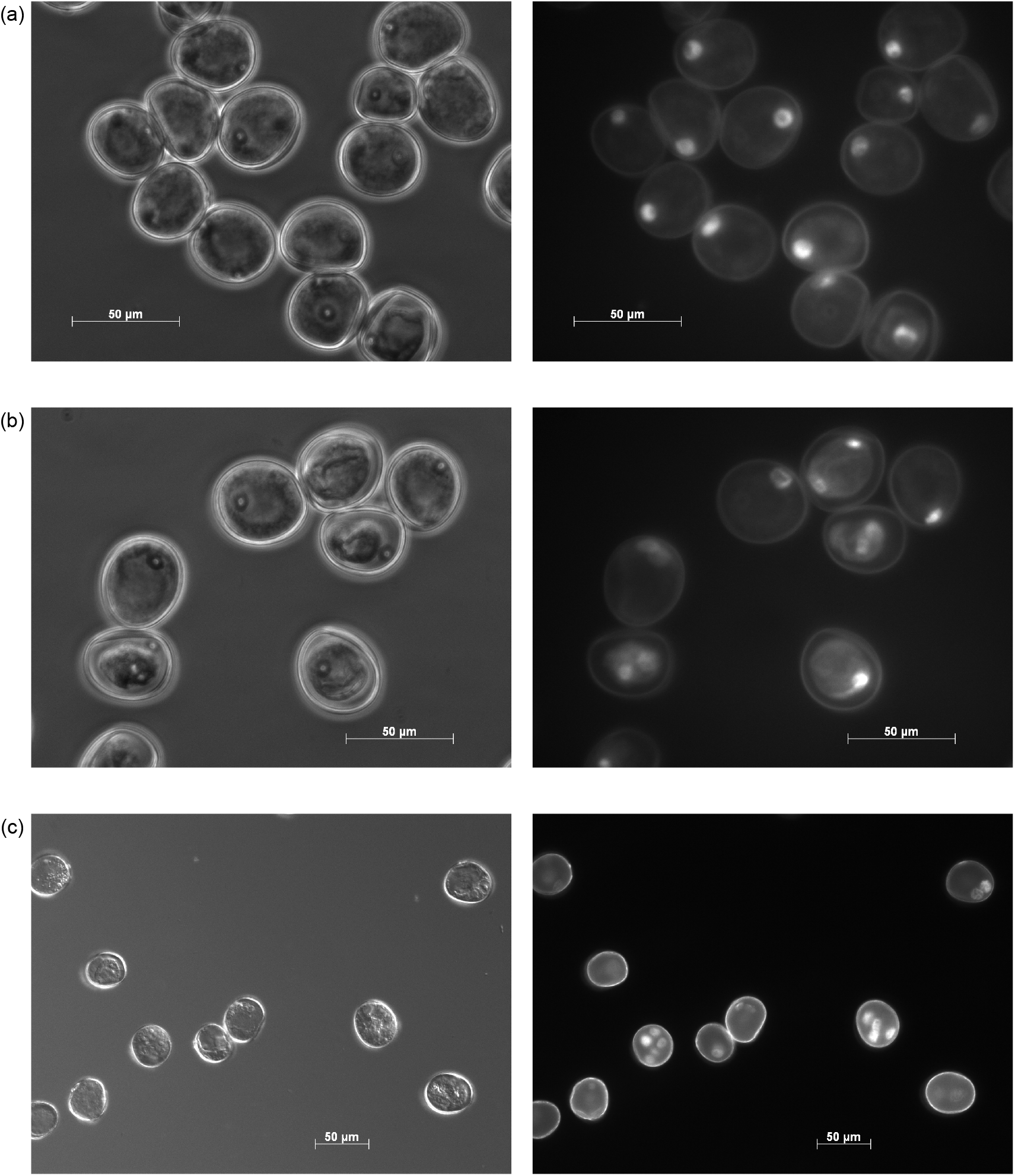
Samples of microspores obtained from the barley cv. Gobernadora at the three stages of early gametic embryogenesis corresponding to (a) freshly harvested microspores without stress (day 0), (b) microspores just after the completion of the stress treatment (day 2) and (c) embryogenesis-induced microspores (day 5). The phenotype of these microspores was captured without (left) and with DAPI staining (right) for microspores on days 0, 2 and 5 using a Zeiss Apoptome microscope under UV laser illumination (excitation of 390/22 nm and emission of 460/50 nm) at a 40x magnification.

### Sequencing sRNAs at three early stages of gametic embryogenesis

We used sRNA-seq to obtain a comprehensive overview of sRNAs expressed in barley microspores at three pivotal time points in the course of gametic embryogenesis. In total, 412.5 million (M) sRNA reads were generated (136.6 M, 128.8 M and 147.1 M reads for microspores on days 0, 2 and 5 respectively) over 12 sRNA libraries (three stages of development x four biological replicates). Reads were trimmed to remove adapters giving 394.8 M clean reads. Of this, a total of 150.1 M (38.0%), 133.1 M (33.7%) and 111.6 M (28.3%) reads, respectively, could be assigned as follows: reads with at least one reported alignment, reads with multiple alignments suppressed due to the -m setting (more than 50 sites), or reads that failed to align to the genome. We considered sRNAs having a read length of 20 to 24-nt as “expressed” if their abundance was equal or higher to 0.5 reads per million (TPM). To assess our experimental repeatability, we performed a multidimensional scaling (MDS) analysis for genome-mapped sRNA reads. We observed three highly distinctive, tight clusters indicating that replicates were uniform and that sRNAs expressed in microspores at each stage (days 0, 2 and 5) were similar (Figure 2). The MDS plot generated from sRNA-seq data was nearly indistinguishable from RNA-seq data previously described by Bélanger et al. (2018). Thus, both the phenotypic (above) and transcriptomic data demonstrated a high degree of experimental reproducibility, thus justifying a joint consideration of these two sources of data (sRNA-seq and previous RNA-seq), to more fully describe and interpret the developmental switch of barley microspores in gametic embryogenesis. In total, 53,189 distinct sRNAs were observed in microspores, of which 7,695, 16,066 and 29,428 sRNAs, respectively, mapped, mapped to multiple sites, or did not map to the genome at all. Until now, sRNAs have not been sequenced in barley microspores. In wheat, Seifert et al. (2016) sequenced and analyzed sRNAs for microspores at similar developmental stages. Despite the fact that we analyzed nearly four times more sequences (394.8 M vs 92.5 M clean reads), we detected fewer sRNAs than Seifert et al. (2016). The lower number of sRNAs detected in barley compared to wheat may be due to the greater genome size (16 Gb vs 5 Gb) and higher ploidy level (6x vs 2x) of the latter. For the ensuing analyses, we focused exclusively on sRNAs that mapped to unique sites in the genome. Among these, the large majority were 24-nt long (6,828; 88.7%) while most of the remaining sRNAs were 21-nt (578; 7.5%), sizes that are typically, but not exclusively, associated mainly with hc-siRNAs and miRNAs. A third category of sRNA, the phasiRNAs, can be either 21-nt or 24-nt. As these three classes of sRNAs are distinct in their biogenesis and functions, we separately categorized, analyzed, and interpreted them as three sRNA groups corresponding to hc-siRNAs, phasiRNAs, and miRNAs.

**Figure 2.**
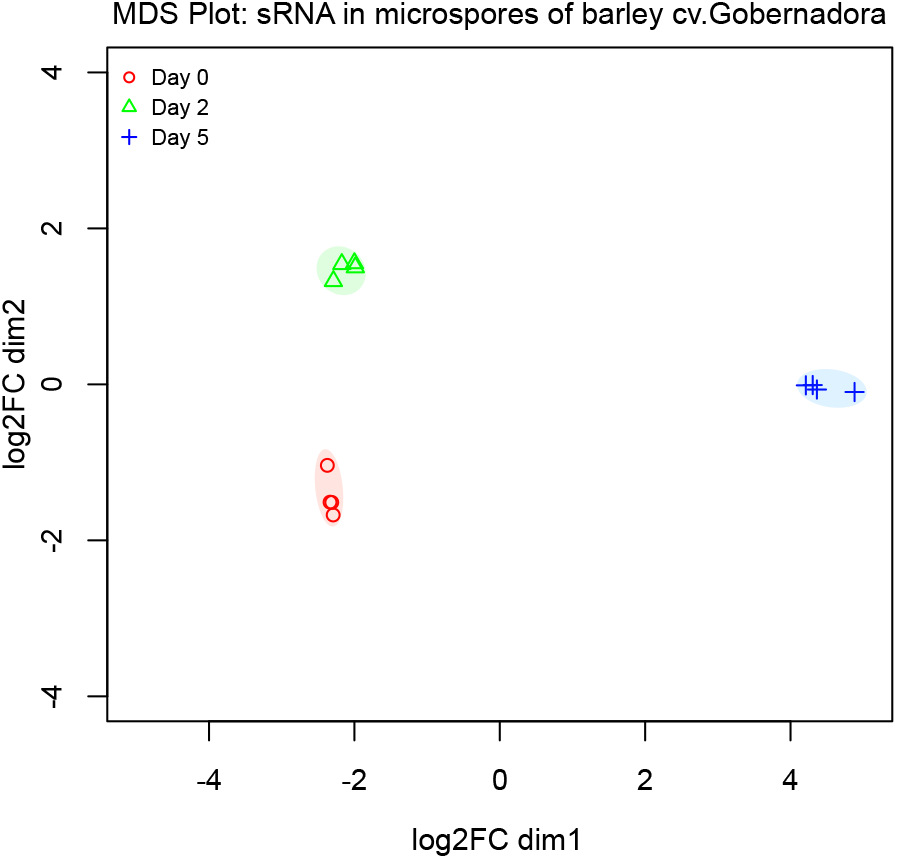
Variation in sRNA expression profiles within and between microspore samples. A MDS plot shows the high degree of uniformity among replicates of the same stage as well as the distinctness between stages.

### There is strict temporal regulation of hc-siRNA throughout microspore development

Known to guide *de novo* DNA methylation of various genomic features (Borges and Martienssen, 2015; Axtell and Meyers, 2018), hc-siRNAs are of interest since they can act in reprogramming microspore genomes. Overall, we detected a total of 6,628 24-nt candidate hc-siRNAs. Of these, the abundance of 3,201 hc-siRNAs (48.3% of all) varied significantly over the course of microspore development at a log2FC ≥ |2.0|. Of these, 879 (13.3%), 1,635 (24.7%) and 3,110 (46.9%) loci were identified when contrasting expression on day 0 vs 2, on day 2 vs 5 as well as on day 0 vs 5, respectively. The visualization of differentially abundant hc-siRNAs revealed three distinct groups corresponding to different expression patterns (Figure 3a). In the first group, the summed abundance of many hc-siRNA loci goes down during the stress treatment applied to microspores (Figure 3a; Cluster 1). In day 2, the abundance of these hc-siRNAs decreased prior to reaching their minimum levels in embryogenic microspores at day 5, suggesting that DNA methylation globally decreased during this developmental period via the RNA-directed DNA methylation (RdDM) pathway. In the research conducted in wheat microspores by Seifert et al. (2016), hc-siRNAs having this type of abundance pattern were not described. In the second group, the summed abundance of many hc-siRNA loci increases during the stress treatment applied to microspores (Figure 3a; Clusters 3 and 4). In a vast majority of cases, the abundance of these hc-siRNAs increased in embryogenic microspores. This suggests an activation of the RdDM pathway for specific genomic features corresponding to these hc-siRNA loci in microspores on day 2, i.e. after completion of the stress treatment. A group of hc-siRNAs having a similar abundance pattern was described by Seifert et al. (2016) in wheat microspores. In the third group, the summed abundance of hundreds of hc-siRNA loci increases markedly during the induction of embryogenesis (Figure 3a; Cluster 2). Most of the hc-siRNAs in this group specifically accumulated in response to the induction of embryogenesis since almost no reads could be detected in microspores on days 0 and 2. Relative to the RdDM pathway, this observation suggests an increase of DNA methylation during the commitment of microspores into embryogenesis. This last group of hc-siRNAs shows an expression pattern consistent with results of Seifert et al. (2016) that described a global increase in 24-nt sRNAs in developing wheat microspores at similar stages of development. Assuming that the abundance of hc-siRNAs and DNA methylation are well correlated, the last group of hc-siRNAs is consistent with observations of El-Tantawy et al. (2014) that measured a progressive increase in DNA methylation from initial microspores to embryos in barley. To summarize, we identified three groups of hc-siRNA exhibiting a strict and distinct temporal regulation. One group drastically decreased in abundance from initial microspores to embryogenic microspores while a second group increased (either abruptly or gradually) in embryogenic microspores. It has been demonstrated that DNA hypomethylation favors microspore reprogramming, totipotency acquisition, and embryogenesis initiation, while embryo differentiation requires *de novo* DNA methylation (Solís et al., 2015; Testillano, 2018). In conclusion, the ability of barley cv. Gobernadora to engage embryonic development might be related to its robust and strict regulation of hc-siRNA in microspores.

**Figure 3.**
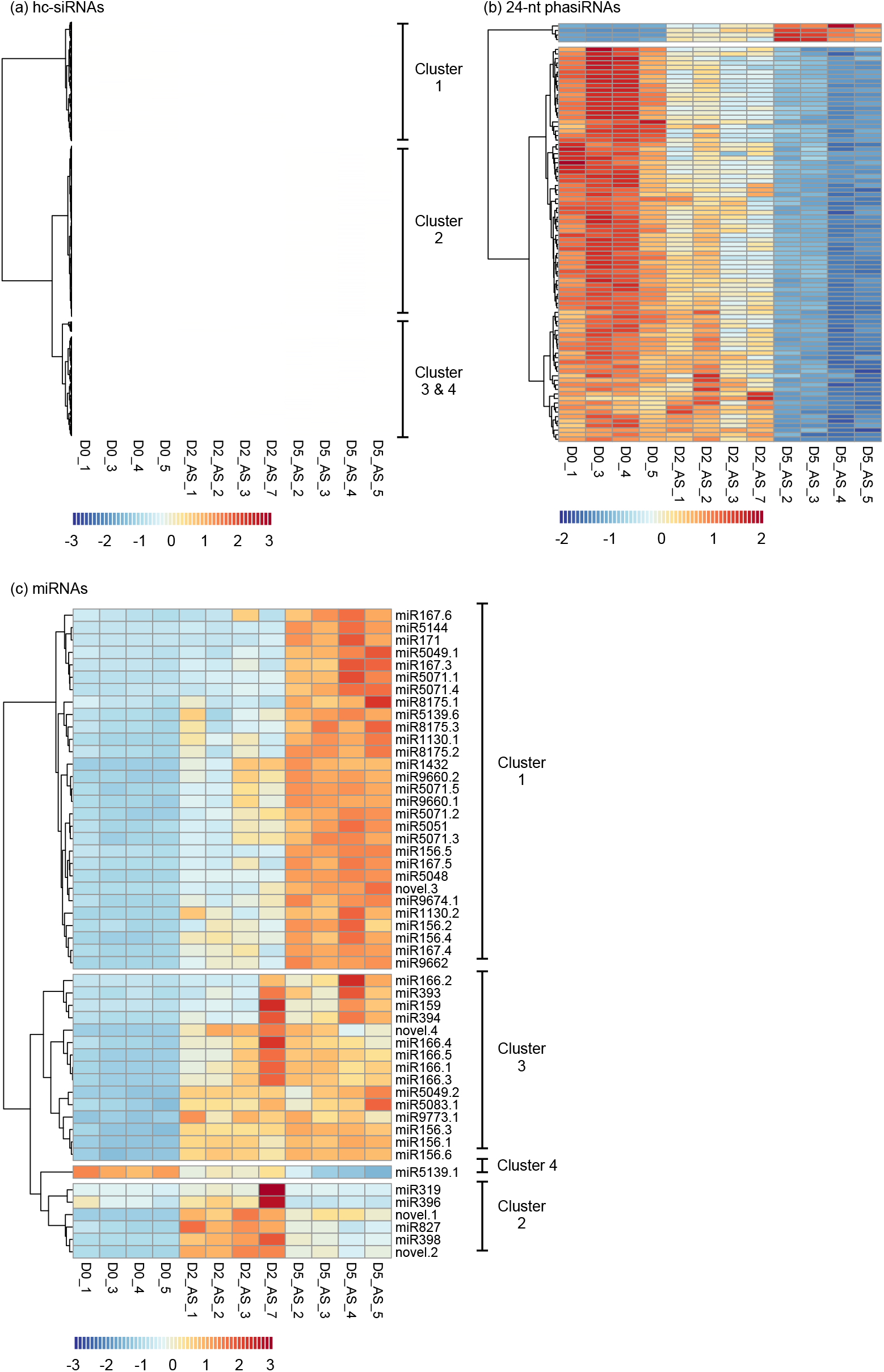
Dynamics of variation in sRNA abundance through barley microspore development. Microspores were analyzed in the early stages of development in androgenesis for differentially expressed (at logFC ≥ |2.0| and *q-value* ≤ 1.0E-03) hc-siRNAs (a) 24-nt phasiRNAs (b) and miRNAs (c). Heatmaps were generated from a read-count matrix normalized for library size and transformed in reads per million. Heatmaps were row normalized.

### 24-nt phasiRNAs regulate protein-coding genes in embryogenic microspores

Among sRNAs expressed in plants, phasiRNAs are a distinct group of sRNAs that are easily distinguishable as a product of processive cleavage of double-stranded RNAs in regular increments (duplexes of 21-nt or 24-nt) from a well-defined terminus (Axtell and Meyers, 2018). To our knowledge, phasiRNAs have never been characterized in the gametic embryogenesis system in barley or that of any other species. Thus, we employed sRNA-seq to annotate phasiRNAs expressed in barley microspores. We detected a total of 202 phasiRNA loci (*PHAS* loci) in barley microspores, almost all of which (192; 95%) originated from 24-*PHAS* loci rather than 21-*PHAS* (10; 5%), loci. All barley *PHAS* loci that we annotated are detailed in Supplementary Table 2. Previous studies describe pre-meiotic (21-*PHAS*) and meiotic (24-*PHAS*) phasiRNA accumulation patterns throughout the development of anthers in maize (Zhai et al., 2015) and rice (Fei et al., 2016). The near absence of 21-nt phasiRNAs that we observed in barley microspores is consistent with the fact that day-0 microspores are post-meiotic cells, and that 21-nt phasiRNAs are produced and accumulate largely in somatic cells (Zhai et al., 2015). Additionally, our observation that all 21-nt phasiRNAs identified in day-0 microspores were significantly repressed in embryogenic day-5 microspores is consistent with the meiotic development patterns reported in the above-mentioned studies in maize and rice. An expression analysis revealed that 90 of the 24-nt barley phasiRNAs that we annotated (46.9% of all 24-nt phasiRNAs) were significantly down-regulated (at log2FC ≥ |2.0|) during the same period (Figure 3b). Moreover, when we relax the threshold to log2FC ≥ |1.5|, the proportion of down-regulated 24-nt phasiRNAs reached 85.4% (164 of 192 24-nt phasiRNAs). When considering that microscopic analysis shows that day-5 microspores are engaged in embryogenesis, a sporophytic development, our observed down-regulation of these phasiRNAs is reasonable, given that 24-nt phasiRNAs have been described as exclusive to reproductive development (Arikit et al., 2013; Zhai et al., 2015; Fei et al., 2016; Yu et al., 2018). Surprisingly, four 24-*PHAS* loci did not follow this general trend, as they were up-regulated in day-5 embryogenic microspores (Figure 3b). Two of these four 24-*PHAS* loci overlapped with protein-coding genes (HORVU1Hr1G006020; HORVU2Hr1G016660), which is not typical of *PHAS* loci, which are commonly derived from long, non-coding RNAs (Komiya, 2017). This observation indicates that the 24-nt phasiRNA pathway can process protein-coding genes after meiosis.

### Up-regulation of miRNAs in embryogenic microspores

In barley, very few miRNAs have been deposited in miRBase and no studies have yet been conducted on microspores; thus, a *de novo* miRNA annotation of barley microspores would provide valuable new information. We annotated a total of 68 miRNAs in microspores, including 62 known and 6 high-confidence novel miRNAs (detailed in Supplementary Tables 3 and 4), according to the criteria for plant miRNA annotation as described by Axtell and Meyers (2018). The 62 known miRNAs covered a total of 30 miRNA families, while each of the new miRNAs represents a candidate novel family. The novel miRNAs we discovered were of relatively low abundance compared to most of the known miRNAs. Thus, it seems likely that our approach to deeply sequence a population of specialized and unique cells in a previously poorly characterized species enabled their discovery. In contrast, Seifert et al. (2016) limited their investigation in wheat to miRNAs already available in miRBase (release 21), excluding known miRNAs conserved with other species, or new miRNA that hadn’t yet been discovered. Additionally, Seifert et al. (2016) limited their description to just four of the 66 miRNAs they found in wheat microspores, consequently, it is not feasible to compare our findings with those of Seifert et al. (2016) in wheat. miRNAs are known to be involved in developmental transitions by targeting families of various transcription factors or other development-related genes (Jones-Rhoades et al., 2006; Chen, 2009; Fei et al., 2016). To identify miRNAs that may play a pivotal role in the developmental switch of microspores into embryogenic development, we performed an expression analysis. We found that the members of four miRNA families accounted for 85.1% of all miRNAs expressed in microspores. Multiple miRNA loci contribute to the abundance of these miRNA family members: miR156, miR166, miR167 and miR9662, whose abundance was estimated at 99 TPM, 251 TPM, 115 TPM and 786 TPM, respectively (Supplementary Table 3). We were unable to find other studies describing these miRNAs, either in gametic or somatic embryogenesis for monocot species closely related to barley. However, Wu et al. (2011) reported that miR156, miR168 and miR171 correlated with the acquisition of embryogenic competence in Valencia sweet orange (a citrus species), as non-embryogenic calli did not express these miRNAs in somatic embryogenesis. A similar finding was described in maize as Juárez-González et al. (2019) showed that miR156 and miR166 accumulate in embryogenic calli. It was further shown that miR156 regulates the expression of *SPL2I4I5I9* thus allowing embryonic competence acquisition in Valencia callus cells (Wu et al., 2011). We detected a significant up-regulation of miR156 expression (6 loci) in microspores from days 0 to 5; thus, it is possible that miR156 may play a pivotal role in barley microspore embryogenesis via the regulation of the *SPL* transcription factor gene family.

We next determined that the abundance of 51 miRNAs (75% of the 68 we annotated) significantly differed between days 0 to 5 of microspore development (detailed in Table 1). In contrast, Seifert et al. (2016) reported that only 4 of 66 miRNAs were differentially expressed in wheat microspores at a similar stage of development. The short (five days total in barley) or long (20 days total in wheat) duration of the stressful treatments applied to microspores might explain the difference in the number of differentially expressed miRNAs between the species. We suggest that sampling wheat microspores over a longer-term treatment may mask expression changes within this interval. That being said, none of the four miRNAs shown to be differentially expressed in wheat were detected in barley microspores, thus we could not make a comparison between the species. In barley, we could distinguish four distinct miRNA expression clusters (Figure 3c; Table 1). One group of miRNAs (Figure 3c; Cluster 2) presented a specific expression pattern: their abundance was strongly induced when the stress treatment was applied to microspores, prior to being repressed when they transitioned to embryogenesis. Among miRNA members of this cluster, we retrieved miR319, miR396, miR398, miR827 and two novel miRNAs (novel.1 and novel.2). The expression pattern of these miRNAs suggests a specific response to the stress treatment applied to microspores. If we consider that stresses applied to microspores will stop pollen grain development, these miRNAs might act by triggering the degradation of key regulators involved in gametic development.

**Table 1.** All differential expression of miRNAs in microspores of barley cv. Gobernadora, contrasted between all three time points (relative to the initiation of the pretreatment inducing the transition towards embryogenesis).

| miRNA | Log2FC for all tested contrasts |  |  |
| --- | --- | --- | --- |
|  | Day 2 - Day 0 | Day 5 - Day 2 | Day 5 - Day 0 |
| **Cluster 1** |  |  |  |
| miR1130.1 | 1.43 | 0.98 | 2.41 |
| miR1130.2 | 3.42 | 0.91 | 4.32 |
| miR1432 | 4.55 | 0.64 | 5.19 |
| miR156.2 | 1.73 | 1.19 | 2.92 |
| miR156.4 | 1.82 | 0.79 | 2.61 |
| miR156.5 | 0.79 | 1.81 | 2.60 |
| miR167.3 | 1.91 | 2.58 | 4.49 |
| miR167.4 | 2.23 | 1.17 | 3.40 |
| miR167.5 | 1.69 | 1.71 | 3.40 |
| miR167.6 | 1.15 | 1.74 | 2.88 |
| miR171 | 1.90 | 4.81 | 6.71 |
| miR5048 | 1.39 | 1.25 | 2.64 |
| miR5049.1 | 0.43 | 2.22 | 2.65 |
| miR5051 | 1.02 | 1.05 | 2.07 |
| miR5071.1 | 1.95 | 2.21 | 4.15 |
| miR5071.2 | 2.25 | 1.09 | 3.34 |
| miR5071.3 | 1.95 | 1.16 | 3.10 |
| miR5071.4 | 1.42 | 2.33 | 3.75 |
| miR5071.5 | 2.18 | 1.16 | 3.33 |
| miR5139.6 | 1.09 | 0.98 | 2.07 |
| miR5144 | 2.12 | 4.26 | 6.39 |
| miR8175.1 | 0.69 | 2.06 | 2.74 |
| miR8175.2 | 1.49 | 1.56 | 3.05 |
| miR8175.3 | 1.45 | 1.48 | 2.92 |
| miR9660.1 | 2.21 | 1.07 | 3.28 |
| miR9660.2 | 2.40 | 0.89 | 3.29 |
| miR9662 | 1.95 | 1.21 | 3.15 |
| miR9674.1 | 1.75 | 1.15 | 2.90 |
| novel.3 | 2.75 | 1.75 | 4.50 |
| **Cluster 2** |  |  |  |
| miR319 | 3.73 | -4.56 | -0.83 |
| miR396 | 1.66 | -3.11 | -1.45 |
| miR398 | 5.85 | -1.52 | 4.33 |
| miR827 | 2.70 | -1.40 | 1.31 |
| novel.1 | 4.08 | -0.73 | 3.35 |
| novel.2 | 4.52 | -1.34 | 3.18 |
| **Cluster 3** |  |  |  |
| miR156.1 | 1.92 | 0.29 | 2.21 |
| miR156.3 | 1.87 | 0.28 | 2.15 |
| miR156.6 | 1.83 | 0.21 | 2.04 |
| miR159 | 3.96 | 0.32 | 4.28 |
| miR166.1 | 2.48 | 0.02 | 2.50 |
| miR166.2 | 2.31 | 1.69 | 4.01 |
| miR166.3 | 2.52 | 0.00 | 2.52 |
| miR166.4 | 2.65 | -0.18 | 2.47 |
| miR166.5 | 2.47 | 0.05 | 2.52 |
| miR393 | 1.82 | 0.73 | 2.55 |
| miR394 | 3.20 | 0.47 | 3.67 |
| miR5049.2 | 2.18 | 0.04 | 2.22 |
| miR5083.1 | 1.99 | 0.13 | 2.11 |
| miR9773.1 | 2.12 | -0.12 | 2.00 |
| novel.4 | 3.11 | -0.40 | 2.71 |
| **Cluster 4** |  |  |  |
| miR5139.1 | -0.74 | -1.65 | -2.39 |

Additional miRNAs were up-regulated in microspores on day 2 after the completion of the stress treatment (Figure 3c; Table 1). The abundance of these miRNAs stayed stable (Figure 3c; Cluster 3) or continued to increase when microspores gained their embryogenic potential on day 5 (Figure 3c; Cluster 1). Among those showing the strongest up-regulation in microspores on day 5 (Table 1), we retrieved miR156, miR159, miR166, miR167, miR171, miR5071, miR1432 and a novel miRNA (novel.3). It seems likely that these miRNAs play a role in the modulation of the stress response, the repression of gametic development, and/or the gain of embryogenic potential. Since the accumulation of all these miRNAs globally increased and stayed high in microspores from day 0 to day 5, they may play a role in the dosage of target transcripts rather than eliminating them altogether. In citrus somatic embryogenesis, concordantly, it was shown that miR156 and miR171 accumulated in embryogenic calli and were correlated with the acquisition of embryogenic competence (Wu et al., 2011). Wu et al. (2011) also reported that miR159 was correlated with the formation of globular-shaped embryos, while miR166 and miR167 were required for cotyledon-shaped embryo morphogenesis. In our biological system, the latter three miRNAs were observed in microspores initiating their first mitotic divisions, a much earlier developmental stage than those described by Wu et al. (2011) in citrus. Although the three latter cases showed a distinction in gametic and somatic embryogenesis systems, more studies are needed to better describe their roles.

### The commitment of microspores into embryogenesis is associated with the miRNA-directed cleavage of ARF, SPL GRF and HD-ZIPIII transcription factor families

To better investigate the role of miRNAs in the developmental switch of microspores, we performed an analysis of miRNA-target interactions. Since transcript cleavage is the principal mechanism of post-transcriptional regulation of miRNAs in plants (Jones-Rhoades et al., 2006; Voinnet, 2009; Shao et al., 2012), we sequenced the 5’ ends of uncapped transcripts to validate miRNA-directed transcript cleavage sites. A total of 141.4 M reads covering one replicate of microspores at the three stages of development (46.8 M, 41.5 M and 53.1 M reads for microspores on days 0, 2 and 5, respectively) was sequenced, mapped to reference protein-coding transcripts and used to validate putative cleavage sites for the microspore-expressed miRNAs we identified in this study (62 known and 6 novel miRNAs). Using these PARE data, we validated a total of 41 distinct miRNA targets in microspores over the three time points. Notably, we were able to validate the cleavage site for one target of the novel.3 miRNA annotated in this study (Table 2; Supplementary Table 5). This result provides strong evidence to confirm our annotation of this novel miRNA. Overall, we validated a total of 8, 25 and 31 miRNA targets in microspores on days 0, 2 and 5, respectively (Supplementary Table 5). We noted that 41.5% of these targets were shared between day-2 and day-5 microspores while 26.8% were unique to day-5 microspores (Table 2). The former set may act to disrupt transcripts involved in pollen development while the latter set may drive the commitment to embryogenesis. Although we deeply sequenced populations of unique cell types, we found the total number of validated targets to be lower than expected. This could be due, in part, to unresolved 5’ UTRs, known as the principal target site of miRNAs, of barley reference transcripts. Alternatively, we don’t know the kinetics of transcript degradation after miRNA-directed cleavage of targeted transcripts; thus, the turnover of cleaved targets might be so rapid that we were unable to capture the uncapped transcripts and validate these miRNA-directed transcript cleavage sites.

**Table 2.**
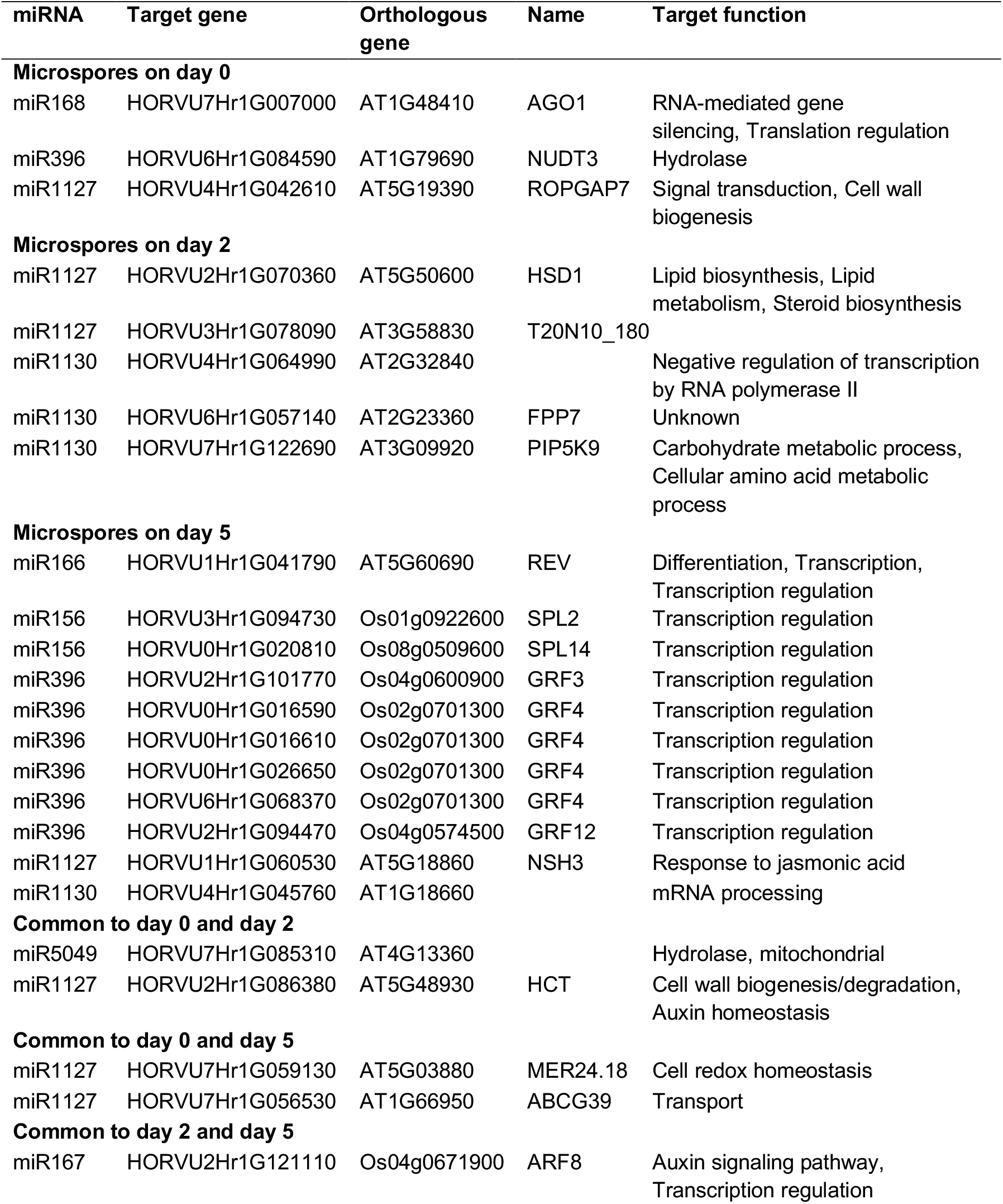
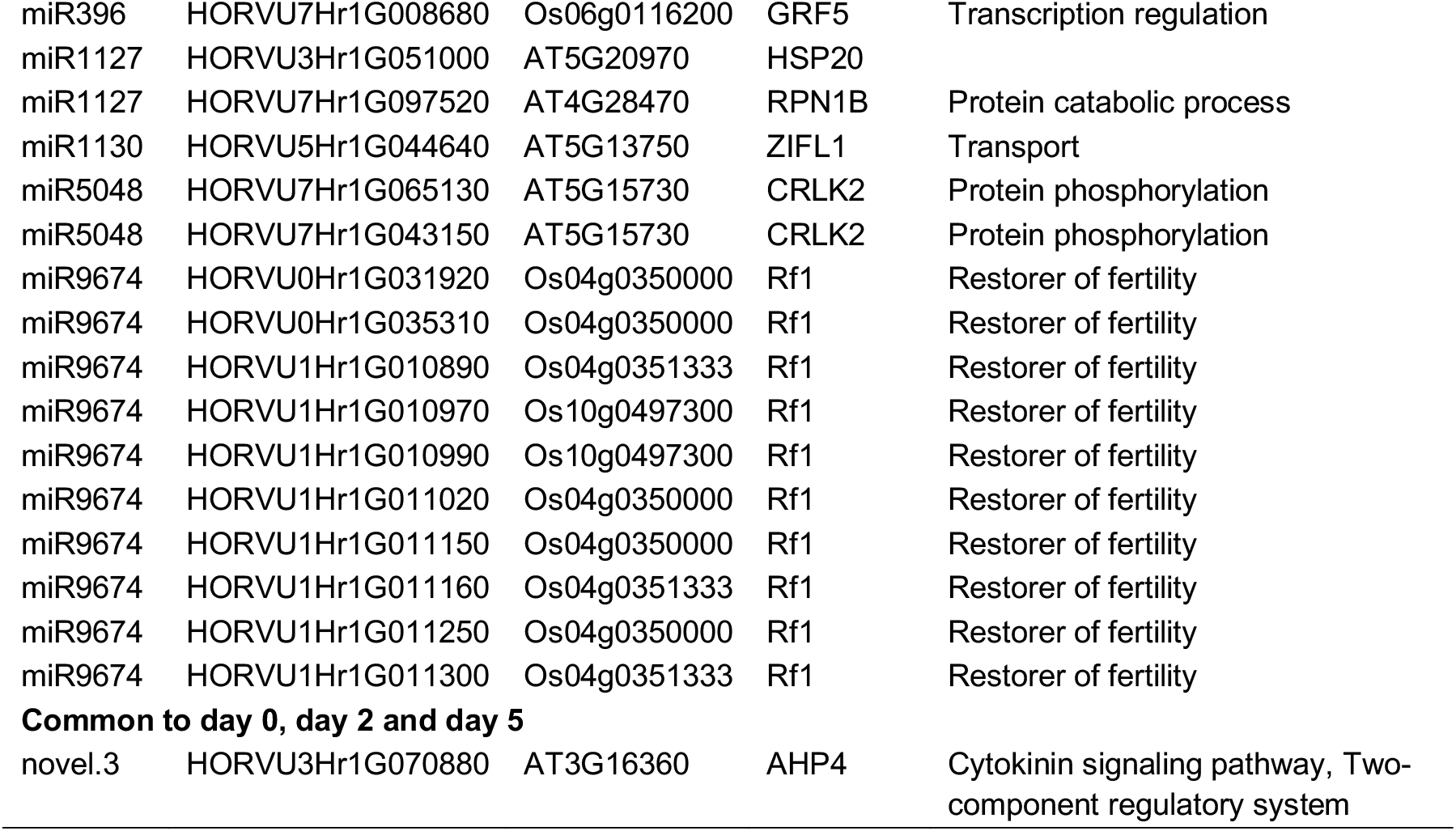
Summary of miRNA targets that were validated via PARE-Seq. The described miRNAs were captured in barley microspores (cv. Gobernadora) at different time points of gametic embyrogenesis.

To better understand the roles that miRNAs may play in the interruption of pollen development, we explored their targets detected in microspores by day 2. In day-2 and day-5 microspores, PARE tags validated the cleavage of HORVU2Hr1G121110 by miR167 (Table 2). This gene is orthologous to *ARF8* in rice. Plant genomes encode several *AUXIN RESPONSE FACTORs* (*ARF*), transcription factors that activate minutes after auxin stimulation, and are involved in various biological developmental processes (Duarte-Aké and De-la-Peña, 2016). Partially redundant, *ARF6* and *ARF8* regulate the maturation of the stamen and the gynoecium by regulating the expression of several developmental genes that coordinate the transition from immature to mature fertile flowers (Nagpal et al., 2005). Loss-of-function mutations in *ARF6* and *ARF8* result in plant sterility (Nagpal et al., 2005). Therefore, our results suggest that miR167 can indirectly interrupt the pollen developmental program via the post-transcriptional regulation of the *ARF8* gene in microspores on day 2. A recent study by Su et al. (2016) showed that over-expression of miR167 in Arabidopsis inhibited somatic embryogenesis; similarly, loss-of-function in *ARF6* and *ARF8* genes resulted in somatic embryogenesis defects. Su et al. (2016) concluded that the miR167-*ARF8* regulation module controls somatic embryogenesis in Arabidopsis. Accordingly, our results suggest that miR167 controls the accumulation of *ARF8* in barley microspores on day 5 and promotes the commitment to embryogenesis.

During the same period of development, we found that miR1130 regulates an ortholog (HORVU5Hr1G044640) of the *ZIFL1* gene (Table 2). A member of the Major Facilitator Superfamily (MFS) transporter in Arabidopsis, the *ZIFL1* gene modulates root auxin-related processes and mediates drought stress tolerance by regulating stomatal closure (Remy et al., 2013). Functional heterologous expression of this plant protein in yeast showed that *ZIFL1* confers increased resistance to auxin-derived herbicides through a reduction of the intracellular concentration of the herbicide 2,4-D (Cabrito et al., 2009). In that study, the herbicide Dicamba was added as a source of auxin to the embryogenesis induction media, suggesting that miR1130 may regulate intracellular auxin concentration through the regulation of the *ZIFL1* gene during the induction of microspore embryogenesis. Additionally, controlling auxin transport and accumulation in microspores may modulate the transcription of *ARF* genes. Thus, we suggest that together, miR1130 and miR167 may act to regulate the concentration of auxin accumulation in microspores, and the subsequent response to this hormonal stimulus.

To study the role that miRNAs can play during the commitment of microspores to embryogenesis, we explored targets exclusive to day-5 microspores. Most of the targets were orthologous to members of the *SQUAMOSA PROMOTER-BINDING-LIKE (SPL), GROWTH-REGULATING FACTOR (GRF)*, and *HD-ZIPIII* transcription factor families. Our results show that miR156 regulates barley orthologs of the rice *SPL2* (HORVU3Hr1G094730) and *SPL14* (HORVU0Hr1G020810) gene in embryogenic microspores (Table 2). Neither *SPL2* or *SPL14* genes have been previously shown to be involved in gametic embryogenesis. However, the miR156-*SPL* regulatory module was inferred to enhance the acquisition of the embryonic competence of citrus calli (Wu et al., 2011; Wu et al., 2015; Long et al., 2018) in somatic embryogenesis by regulating *SPL2I4I5I9*. Also, we observed that miR396 regulates several members of the *GRF* family of genes (Table 2), namely *GRF3* (HORVU2Hr1G101770), *GRF4* (HORVU0Hr1G016590; HORVU0Hr1G016610; HORVU0Hr1G026650; HORVU6Hr1G068370), *GRF5* (HORVU7Hr1G008680) and *GRF12* (HORVU2Hr1G094470). Many plant genomes encode several *GRF* proteins known to be involved in multiple developmental processes (Liebsch and Palatnik, 2020). In rice, *GRF4* has been described as a positive regulator of genes that promote cell proliferation (Hu et al., 2015; Sun et al., 2016) and the miR396-*GRF* module may regulate cell proliferation in meristematic tissues. Furthermore, rice *GRF4* activates transcription of expansin promoters in protoplasts suggesting a potential function in cell expansion (Liebsch and Palatnik, 2020). We speculate that miR396 controls the accumulation of *GRF* genes to facilitate sufficient mitotic divisions in microspores to generate a multicellular structure prior to the activation of the embryo maturation program. While none of the miR396-*GRF* interactions that we identified have previously been implicated in gametic embryogenesis, Szczygieł-Sommer and Ga (2019) showed recently that the induction of somatic embryogenesis was enhanced by the regulation of *GRF1, GRF4, GRF7, GRF8* and *GRF9* genes by miR396; this contributed to controlling the sensitivity of tissues to auxin treatment and enhanced embryogenesis induction in Arabidopsis. Finally, we found that miR166 targeted a barley ortholog to the Arabidopsis *REVOLUTA* gene (*REV*; HORVU1Hr1G041790) (Table 2). In Arabidopsis, PHABULOSA (PHB), PHAVOLUTA (PHV) and REV are all members of the HD-ZIPIII transcription factor family, and are regulators of early zygotic embryo development (Nowak and Gaj, 2016). As proposed by Wu et al. (2015), the miR166-*REV* hub may prevent precocious activation of the embryo maturation program in citrus somatic embryogenesis. Although the roles of *SPL*, *GRF* and REV are likely distinct during induction of embryogenesis, the regulatory activity of miR156-*SPL*, miR396-*GRF* and miR166-*REV* modules in gametic or somatic embryogenesis in eudicot or monocot species suggests a functional conservation of these regulatory hubs during the acquisition of embryogenic potential for both systems.

## CONCLUDING REMARKS

Our work, and that of Bélanger et al. (2018), contribute a database of gene expression (RNA, sRNA and PARE sequencing) in microspore development during embryogenesis. We chose to use the barley cultivar Gobernadora due to its high propensity for microspore conversion to embryos, despite the fact that Gobernadora is highly susceptible to albinism. Previous gametic embryogenesis studies were performed on distinct species, varieties for a given species, and using diverse experimental approaches such as various stress pretreatments and embryogenesis media formulation, thus limiting our ability to compare studies and to identify molecular determinants of microspore development. We hope that additional studies using the same barley cultivar (cv. Gobernadora), pretreatments, and culture protocols (isolated microspores) will be used for various approaches, including genomics, epigenetics, transcriptomics or proteomics. Such coordinated efforts will contribute to the development of a model for molecular changes in plant cell fate decisions.

## AUTHOR CONTRIBUTIONS

S.B. and F.B designed the research; S.M. and P.E. prepared plant material; P.B. constructed sRNA and PARE libraries; S.B analyzed and interpreted the data; M.A.L. contributed to PARE analysis; P.E.J., B.C.M and F.B. contributed to data interpretation; S.B. wrote the manuscript; B.C.M and F.B. reviewed the manuscript.

The author responsible for distribution of materials integral to the findings presented in this article is: François Belzile

## ACKNOWLEDGMENTS

We thank Joanna Friesner for assistance with editing. S. Bélanger gratefully acknowledges graduate studentships from the National Sciences and Engineering Research Council of Canada (NSERC) and the NSERC CREATE AgroPhytoSciences program. This work was supported by a research grant from the NSERC of Canada to F. Belzile, as well as resources from the Université Laval – Québec. This research was enabled in part by support provided by Calcul Québec and Compute Canada.

---

**Supplementary Table 1.**
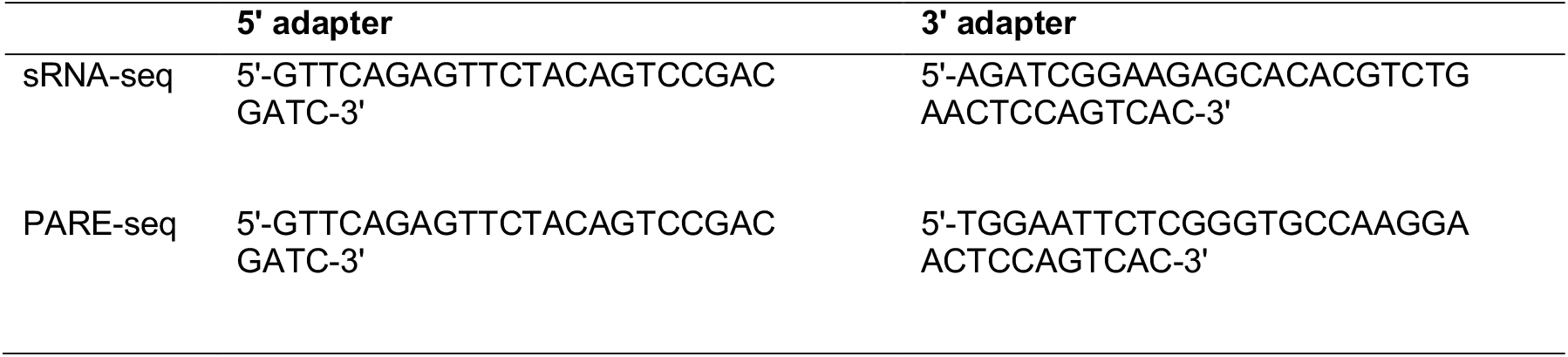
The 5’ and 3’ adapters used to construct libraries.

**Supplementary Table 2.** Coordinates and abundance of the 202 PHAS loci detected in barley microspores (cv. Gobernadora) undergoing gametic embyrogenesis.

| *PHAS* loci ID | Chr | Start | End | Total abundance captured over 12<br>microspore libraries (rpm) |
| --- | --- | --- | --- | --- |
| 21PHAS_14322 | 3H | 11468203 | 11468794 | 21.3 |
| 21PHAS_20868 | 3H | 666495064 | 666495851 | 3.8 |
| 21PHAS_20987 | 3H | 671295658 | 671296326 | 2.7 |
| 21PHAS_33492 | 5H | 584990830 | 584991582 | 1.9 |
| 21PHAS_34003 | 5H | 610464867 | 610465221 | 1.5 |
| 21PHAS_35755 | 6H | 15137173 | 15137811 | 2.4 |
| 21PHAS_42227 | 7H | 21074671 | 21075109 | 3.6 |
| 21PHAS_45781 | 7H | 414260885 | 414264622 | 8.1 |
| 21PHAS_47766 | 7H | 614055096 | 614055820 | 1.8 |
| 21PHAS_41814 | 7H | 7355310 | 7355944 | 4.6 |
| 24PHAS_263 | 1H | 13054359 | 13054962 | 25.6 |
| 24PHAS_268 | 1H | 13068606 | 13070067 | 107.3 |
| 24PHAS_272 | 1H | 13081265 | 13081853 | 39.6 |
| 24PHAS_46 | 1H | 2112051 | 2112791 | 10.0 |
| 24PHAS_452 | 1H | 21184712 | 21185479 | 20.8 |
| 24PHAS_472 | 1H | 21635818 | 21636352 | 6.5 |
| 24PHAS_478 | 1H | 21906855 | 21907639 | 5.6 |
| 24PHAS_480 | 1H | 21912182 | 21912444 | 2.9 |
| 24PHAS_484 | 1H | 22024537 | 22025274 | 12.0 |
| 24PHAS_492 | 1H | 22164473 | 22165032 | 10.8 |
| 24PHAS_3397 | 1H | 364128661 | 364132480 | 36.7 |
| 24PHAS_104 | 1H | 4694003 | 4694589 | 9.4 |
| 24PHAS_4720 | 1H | 487210157 | 487210328 | 58.9 |
| 24PHAS_4723 | 1H | 487322711 | 487323442 | 253.0 |
| 24PHAS_112 | 1H | 5141100 | 5142006 | 14.0 |
| 24PHAS_5591 | 1H | 531451194 | 531452501 | 1.7 |
| 24PHAS_6022 | 1H | 544721420 | 544721891 | 5.0 |
| 24PHAS_1120 | 1H | 74270142 | 74270793 | 15.2 |
| 24PHAS_6878 | 2H | 16876358 | 16877406 | 8.2 |
| 24PHAS_6891 | 2H | 17054345 | 17055506 | 4.5 |
| 24PHAS_6895 | 2H | 17089892 | 17090399 | 5.7 |
| 24PHAS_6901 | 2H | 17158984 | 17159895 | 3.0 |
| 24PHAS_9080 | 2H | 188226877 | 188228839 | 331.8 |
| 24PHAS_6992 | 2H | 19237294 | 19238107 | 11.4 |
| 24PHAS_7004 | 2H | 19326424 | 19327410 | 3.4 |
| 24PHAS_7009 | 2H | 19367354 | 19368148 | 5.6 |
| 24PHAS_7013 | 2H | 19463512 | 19464618 | 8.7 |
| 24PHAS_7025 | 2H | 19689880 | 19690467 | 7.1 |
| 24PHAS_7152 | 2H | 24564837 | 24565433 | 18.6 |
| 24PHAS_7162 | 2H | 24899805 | 24900585 | 21.0 |
| 24PHAS_7265 | 2H | 29567069 | 29568231 | 4.0 |
| 24PHAS_7474 | 2H | 38376025 | 38378231 | 182.6 |
| 24PHAS_11697 | 2H | 586819680 | 586822269 | 248.9 |
| 24PHAS_12394 | 2H | 663464067 | 663466596 | 53.4 |
| 24PHAS_6621 | 2H | 6708667 | 6709785 | 2.8 |
| 24PHAS_12564 | 2H | 679346270 | 679347678 | 2.4 |
| 24PHAS_6627 | 2H | 6991738 | 6992043 | 5.8 |
| 24PHAS_13189 | 2H | 715835306 | 715835812 | 72.0 |
| 24PHAS_13281 | 2H | 717666992 | 717667913 | 5.5 |
| 24PHAS_13462 | 2H | 728781631 | 728782617 | 3.8 |
| 24PHAS_13482 | 2H | 729376598 | 729377279 | 5.5 |
| 24PHAS_13483 | 2H | 729400706 | 729401490 | 42.0 |
| 24PHAS_13486 | 2H | 729497742 | 729498130 | 8.7 |
| 24PHAS_13489 | 2H | 729558100 | 729558949 | 7.4 |
| 24PHAS_13685 | 2H | 744773597 | 744774368 | 6.5 |
| 24PHAS_13804 | 2H | 751202233 | 751203753 | 20.0 |
| 24PHAS_13852 | 2H | 753195658 | 753196020 | 3.9 |
| 24PHAS_13853 | 2H | 753200440 | 753201076 | 1.6 |
| 24PHAS_13895 | 2H | 755104957 | 755106046 | 7.1 |
| 24PHAS_13986 | 2H | 760732391 | 760733419 | 1.7 |
| 24PHAS_14019 | 2H | 762094431 | 762095616 | 1.7 |
| 24PHAS_14020 | 2H | 762116991 | 762118168 | 1.8 |
| 24PHAS_14093 | 2H | 764844581 | 764845875 | 13.5 |
| 24PHAS_14108 | 2H | 765730662 | 765732083 | 251.3 |
| 24PHAS_14110 | 2H | 765919784 | 765920827 | 4.8 |
| 24PHAS_14353 | 3H | 11740486 | 11741320 | 15.4 |
| 24PHAS_14383 | 3H | 12085831 | 12086703 | 4.2 |
| 24PHAS_14384 | 3H | 12120475 | 12121345 | 14.1 |
| 24PHAS_14385 | 3H | 12123138 | 12123852 | 7.4 |
| 24PHAS_14391 | 3H | 12294741 | 12295148 | 2.3 |
| 24PHAS_14392 | 3H | 12296082 | 12297280 | 3.5 |
| 24PHAS_14474 | 3H | 16256234 | 16257111 | 2.8 |
| 24PHAS_16117 | 3H | 185045546 | 185047175 | 102.2 |
| 24PHAS_14534 | 3H | 18905936 | 18908130 | 3.6 |
| 24PHAS_14698 | 3H | 26975711 | 26976984 | 3.7 |
| 24PHAS_17125 | 3H | 314356585 | 314358472 | 27.8 |
| 24PHAS_14205 | 3H | 3661188 | 3662229 | 5.8 |
| 24PHAS_18743 | 3H | 525824091 | 525824462 | 42.1 |
| 24PHAS_15179 | 3H | 63340143 | 63340569 | 74.9 |
| 24PHAS_20715 | 3H | 664150651 | 664151323 | 4.3 |
| 24PHAS_20725 | 3H | 664327397 | 664327804 | 4.5 |
| 24PHAS_15208 | 3H | 66434284 | 66436103 | 3.0 |
| 24PHAS_20728 | 3H | 664359557 | 664360104 | 3.7 |
| 24PHAS_20911 | 3H | 667876276 | 667877972 | 6.7 |
| 24PHAS_21215 | 3H | 683875004 | 683877450 | 2.3 |
| 24PHAS_21219 | 3H | 683899084 | 683899910 | 21.1 |
| 24PHAS_21221 | 3H | 683902316 | 683903381 | 8.1 |
| 24PHAS_21233 | 3H | 684159299 | 684160072 | 58.9 |
| 24PHAS_21235 | 3H | 684171527 | 684173251 | 23.6 |
| 24PHAS_21240 | 3H | 684233486 | 684234316 | 3.2 |
| 24PHAS_21242 | 3H | 684242678 | 684243510 | 6.1 |
| 24PHAS_21248 | 3H | 684372625 | 684373460 | 17.5 |
| 24PHAS_21251 | 3H | 684381656 | 684382642 | 9.8 |
| 24PHAS_21254 | 3H | 684389081 | 684390158 | 9.8 |
| 24PHAS_21574 | 3H | 698405608 | 698406032 | 19.1 |
| 24PHAS_14254 | 3H | 7693336 | 7695103 | 2.4 |
| 24PHAS_21614 | 4H | 1232114 | 1233156 | 5.8 |
| 24PHAS_21886 | 4H | 12435409 | 12438601 | 70.7 |
| 24PHAS_23512 | 4H | 169628030 | 169631418 | 95.3 |
| 24PHAS_22324 | 4H | 37361049 | 37362224 | 15.0 |
| 24PHAS_22326 | 4H | 37379486 | 37380118 | 9.8 |
| 24PHAS_22338 | 4H | 37681972 | 37682964 | 40.4 |
| 24PHAS_22339 | 4H | 37695386 | 37696271 | 10.1 |
| 24PHAS_22388 | 4H | 43171499 | 43172430 | 43.0 |
| 24PHAS_22389 | 4H | 43181935 | 43182742 | 30.1 |
| 24PHAS_22392 | 4H | 43493799 | 43494612 | 32.7 |
| 24PHAS_27704 | 4H | 622935559 | 622936823 | 15.8 |
| 24PHAS_27912 | 4H | 633812349 | 633814825 | 24.9 |
| 24PHAS_29466 | 5H | 109616707 | 109620398 | 39.5 |
| 24PHAS_31814 | 5H | 453066126 | 453072802 | 228.6 |
| 24PHAS_32291 | 5H | 507349064 | 507349976 | 23.9 |
| 24PHAS_32774 | 5H | 548843429 | 548845358 | 2.4 |
| 24PHAS_32879 | 5H | 553897889 | 553898723 | 12.4 |
| 24PHAS_32880 | 5H | 553971675 | 553972443 | 3.2 |
| 24PHAS_32970 | 5H | 556833630 | 556834202 | 10.4 |
| 24PHAS_28273 | 5H | 5587310 | 5588002 | 39.9 |
| 24PHAS_28274 | 5H | 5588957 | 5589947 | 6.0 |
| 24PHAS_33457 | 5H | 582637654 | 582639849 | 7.3 |
| 24PHAS_34293 | 5H | 623928857 | 623930166 | 2.9 |
| 24PHAS_34365 | 5H | 626025368 | 626027216 | 182.2 |
| 24PHAS_28314 | 5H | 6319371 | 6320356 | 13.4 |
| 24PHAS_34521 | 5H | 632530990 | 632531931 | 25.6 |
| 24PHAS_28317 | 5H | 6341068 | 6342316 | 22.5 |
| 24PHAS_28320 | 5H | 6352728 | 6353329 | 14.7 |
| 24PHAS_28323 | 5H | 6372769 | 6373300 | 3.8 |
| 24PHAS_28331 | 5H | 6499566 | 6500489 | 8.4 |
| 24PHAS_34940 | 5H | 652041234 | 652043511 | 2.8 |
| 24PHAS_28334 | 5H | 6551607 | 6553132 | 17.5 |
| 24PHAS_35108 | 5H | 659336027 | 659337316 | 7.3 |
| 24PHAS_35149 | 5H | 660944471 | 660945690 | 5.2 |
| 24PHAS_35160 | 5H | 661060785 | 661063176 | 4.0 |
| 24PHAS_29363 | 5H | 94270838 | 94271647 | 45.9 |
| 24PHAS_35606 | 6H | 10954184 | 10954730 | 5.6 |
| 24PHAS_35608 | 6H | 10956100 | 10956941 | 21.0 |
| 24PHAS_35610 | 6H | 10960288 | 10960790 | 18.2 |
| 24PHAS_35614 | 6H | 11071879 | 11072528 | 15.1 |
| 24PHAS_35615 | 6H | 11073419 | 11074092 | 29.8 |
| 24PHAS_35618 | 6H | 11088920 | 11089614 | 44.1 |
| 24PHAS_35622 | 6H | 11123413 | 11124257 | 23.4 |
| 24PHAS_35628 | 6H | 11301466 | 11302319 | 8.8 |
| 24PHAS_35806 | 6H | 16645266 | 16646341 | 2.4 |
| 24PHAS_36128 | 6H | 36173676 | 36174384 | 6.6 |
| 24PHAS_36266 | 6H | 44042823 | 44047172 | 97.6 |
| 24PHAS_40746 | 6H | 543603160 | 543603685 | 4.8 |
| 24PHAS_40748 | 6H | 543700616 | 543701081 | 14.7 |
| 24PHAS_40749 | 6H | 543734455 | 543735267 | 21.3 |
| 24PHAS_40750 | 6H | 543737752 | 543738859 | 6.4 |
| 24PHAS_40751 | 6H | 543856342 | 543857069 | 7.2 |
| 24PHAS_40752 | 6H | 543858757 | 543859066 | 4.7 |
| 24PHAS_40957 | 6H | 555942455 | 555943569 | 2.5 |
| 24PHAS_41280 | 6H | 570939408 | 570942261 | 6.7 |
| 24PHAS_41285 | 6H | 571024040 | 571025511 | 18.7 |
| 24PHAS_41286 | 6H | 571027635 | 571028320 | 2.7 |
| 24PHAS_41289 | 6H | 571041270 | 571041644 | 84.6 |
| 24PHAS_36607 | 6H | 70476310 | 70477145 | 19.1 |
| 24PHAS_41963 | 7H | 11341442 | 11342513 | 3.9 |
| 24PHAS_41964 | 7H | 11380125 | 11381193 | 3.8 |
| 24PHAS_42015 | 7H | 14297407 | 14297945 | 13.7 |
| 24PHAS_42017 | 7H | 14419777 | 14419974 | 10.7 |
| 24PHAS_42019 | 7H | 14435981 | 14436570 | 12.5 |
| 24PHAS_42021 | 7H | 14447177 | 14448617 | 5.3 |
| 24PHAS_42023 | 7H | 14484536 | 14485400 | 4.0 |
| 24PHAS_42030 | 7H | 14735397 | 14736383 | 11.8 |
| 24PHAS_42041 | 7H | 14949791 | 14950623 | 35.7 |
| 24PHAS_42052 | 7H | 15502548 | 15502916 | 2.3 |
| 24PHAS_42055 | 7H | 15527552 | 15528404 | 4.0 |
| 24PHAS_42057 | 7H | 15543784 | 15544259 | 9.6 |
| 24PHAS_42059 | 7H | 15574368 | 15574914 | 11.2 |
| 24PHAS_42063 | 7H | 15617409 | 15617717 | 12.4 |
| 24PHAS_42075 | 7H | 15667079 | 15667886 | 9.5 |
| 24PHAS_42078 | 7H | 15673681 | 15674595 | 71.8 |
| 24PHAS_42092 | 7H | 15934967 | 15935551 | 4.1 |
| 24PHAS_42099 | 7H | 16188506 | 16188795 | 6.0 |
| 24PHAS_42100 | 7H | 16188973 | 16189373 | 4.9 |
| 24PHAS_42470 | 7H | 32268202 | 32269098 | 4.4 |
| 24PHAS_42617 | 7H | 41001346 | 41002354 | 23.0 |
| 24PHAS_42619 | 7H | 41005367 | 41006409 | 8.7 |
| 24PHAS_42653 | 7H | 43887144 | 43888996 | 17.3 |
| 24PHAS_42711 | 7H | 46466089 | 46467935 | 4.7 |
| 24PHAS_47112 | 7H | 566467182 | 566467765 | 9.0 |
| 24PHAS_47139 | 7H | 568548901 | 568549595 | 11.3 |
| 24PHAS_47146 | 7H | 568689134 | 568690021 | 3.7 |
| 24PHAS_47187 | 7H | 571782434 | 571782975 | 5.5 |
| 24PHAS_47188 | 7H | 571792628 | 571793469 | 30.9 |
| 24PHAS_47889 | 7H | 619383864 | 619384720 | 130.7 |
| 24PHAS_48004 | 7H | 625274352 | 625274927 | 1.8 |
| 24PHAS_48144 | 7H | 632308011 | 632308901 | 3.2 |
| 24PHAS_48157 | 7H | 633013157 | 633014228 | 3.7 |
| 24PHAS_48305 | 7H | 636391193 | 636392080 | 2.6 |
| 24PHAS_48306 | 7H | 636408765 | 636409414 | 3.9 |
| 24PHAS_48312 | 7H | 636582711 | 636584183 | 5.1 |
| 24PHAS_48337 | 7H | 637600962 | 637601806 | 9.5 |
| 24PHAS_48343 | 7H | 637666684 | 637667402 | 21.3 |
| 24PHAS_48345 | 7H | 637729899 | 637730788 | 11.8 |
| 24PHAS_48551 | 7H | 643000930 | 643002647 | 5.5 |
| 24PHAS_48650 | 7H | 645403322 | 645404928 | 4.0 |
| 24PHAS_48780 | 7H | 649918353 | 649919340 | 4.3 |
| 24PHAS_48870 | 7H | 652224124 | 652224805 | 7.2 |
| 24PHAS_49727 | Un | 101568653 | 101572067 | 14.6 |
| 24PHAS_49094 | Un | 9394321 | 9394968 | 2.3 |
| 24PHAS_49097 | Un | 9401319 | 9402253 | 4.5 |
| 24PHAS_49099 | Un | 9418578 | 9419171 | 5.9 |

**Supplementary Table 3.** Coordinates and abundance of the 68 annotated miRNAs in barley microspores (cv. Gobernadora) undergoing gametic embyrogenesis.

| miRNA | Chromosome | Start | End | Total abundance captured<br>over 12 microspore libraries<br>(rpm) |
| --- | --- | --- | --- | --- |
| miR1127 | 2H | 156304849 | 156306165 | 1.4 |
| miR1130.1 | 1H | 517537524 | 517539284 | 10.0 |
| miR1130.2 | 5H | 558876253 | 558877962 | 0.7 |
| miR1432 | 2H | 166771786 | 166771953 | 0.9 |
| miR156.1 | 2H | 623451730 | 623451867 | 14.8 |
| miR156.2 | 3H | 59655592 | 59655682 | 0.9 |
| miR156.3 | 3H | 59655786 | 59655875 | 7.0 |
| miR156.4 | 5H | 499573181 | 499573663 | 5.1 |
| miR156.5 | 5H | 499573813 | 499574078 | 2.4 |
| miR156.6 | 6H | 362942108 | 362943254 | 68.4 |
| miR159 | 3H | 14332438 | 14332747 | 24.6 |
| miR166.1 | 1H | 353507033 | 353507137 | 92.1 |
| miR166.2 | 5H | 513807917 | 513808017 | 2.5 |
| miR166.3 | 5H | 593550048 | 593550127 | 84.7 |
| miR166.4 | 6H | 539379081 | 539379103 | 1.6 |
| miR166.5 | 6H | 539379309 | 539379352 | 29.5 |
| miR166.6 | 7H | 558749092 | 558749512 | 40.6 |
| miR167.1 | 4H | 590940939 | 590941052 | 1.3 |
| miR167.2 | 4H | 591009157 | 591009266 | 1.4 |
| miR167.3 | 5H | 595203284 | 595203376 | 1.9 |
| miR167.4 | 6H | 136921506 | 136921623 | 5.9 |
| miR167.5 | Un | 58632427 | 58632921 | 103.9 |
| miR167.6 | Un | 58635472 | 58635548 | 0.5 |
| miR168 | 6H | 34582524 | 34582666 | 27.4 |
| miR171 | 4H | 617878928 | 617879096 | 1.5 |
| miR1878 | 5H | 479890906 | 479890931 | 2.0 |
| miR319 | 3H | 140433531 | 140433705 | 3.0 |
| miR393 | 2H | 756463338 | 756463486 | 0.6 |
| miR394 | 3H | 649738003 | 649739841 | 0.6 |
| miR396 | 6H | 564855150 | 564855213 | 1.1 |
| miR398 | 3H | 637805239 | 637806060 | 1.7 |
| miR408 | 7H | 622016838 | 622016929 | 0.6 |
| miR444 | 2H | 583382959 | 583383500 | 2.3 |
| miR5048 | 7H | 550741190 | 550742460 | 28.9 |
| miR5049.1 | 3H | 95702904 | 95703500 | 1.0 |
| miR5049.2 | 1H | 534280976 | 534281606 | 1.3 |
| miR5051 | 4H | 588904352 | 588906553 | 7.0 |
| miR5071.1 | 3H | 667572215 | 667572472 | 6.8 |
| miR5071.2 | 3H | 667603611 | 667603714 | 1.2 |
| miR5071.3 | 4H | 387684886 | 387685014 | 1.7 |
| miR5071.4 | 5H | 289162901 | 289163053 | 0.6 |
| miR5071.5 | 6H | 572661234 | 572661255 | 1.9 |
| miR5083.1 | 1H | 554520362 | 554520768 | 1.2 |
| miR5083.2 | 1H | 554613391 | 554613666 | 0.6 |
| miR5139.1 | 2H | 457339091 | 457339814 | 0.6 |
| miR5139.2 | 3H | 267694291 | 267694389 | 2.7 |
| miR5139.3 | 3H | 546431702 | 546431746 | 0.9 |
| miR5139.4 | 5H | 325277882 | 325277997 | 1.5 |
| miR5139.5 | 7H | 566782938 | 566783769 | 1.0 |
| miR5139.6 | Un | 126464383 | 126464540 | 6.0 |
| miR5144 | 2H | 23937585 | 23939427 | 1.3 |
| miR8175.1 | 4H | 285327304 | 285327355 | 0.6 |
| miR8175.2 | 4H | 611607026 | 611607134 | 2.9 |
| miR8175.3 | 7H | 164139781 | 164139848 | 1.3 |
| miR827 | 2H | 620316184 | 620316763 | 3.9 |
| miR9660.1 | 6H | 558257705 | 558257876 | 4.3 |
| miR9660.2 | 6H | 558398762 | 558398935 | 4.5 |
| miR9662 | 6H | 104007160 | 104007445 | 735.9 |
| miR9674.1 | 3H | 638444020 | 638444191 | 3.0 |
| miR9674.2 | 6H | 15557530 | 15557700 | 13.4 |
| miR9773.1 | 3H | 673282151 | 673282417 | 16.6 |
| miR9773.2 | 7H | 329970778 | 329971374 | 0.7 |
| novel.1 | 1H | 6577445 | 6577556 | 1.0 |
| novel.2 | 4H | 192456595 | 192456819 | 1.9 |
| novel.3 | 5H | 471378581 | 471378848 | 0.7 |
| novel.4 | 7H | 532917020 | 532917319 | 4.7 |
| novel.5 | 7H | 575649579 | 575649830 | 5.7 |
| novel.6 | 7H | 593828495 | 593828636 | 0.9 |

**Supplementary Table 4.**
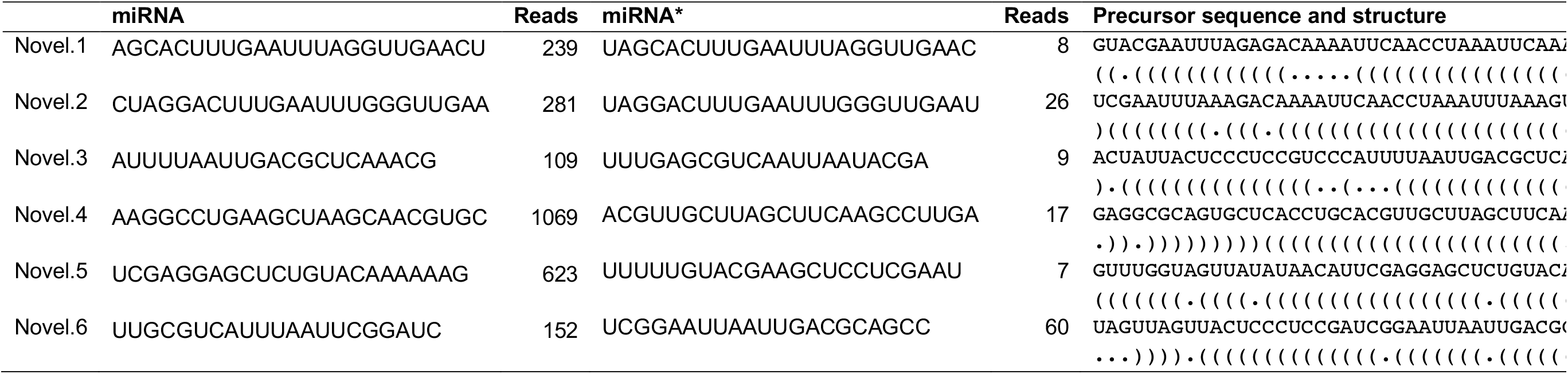
Sequences of mature miRNA, miRNA* and their putative precursors as well as their abundance detected in microspores of barley cv. Gobernadora undergoing gametic embyrogenesis.

**Supplementary Table 5.**
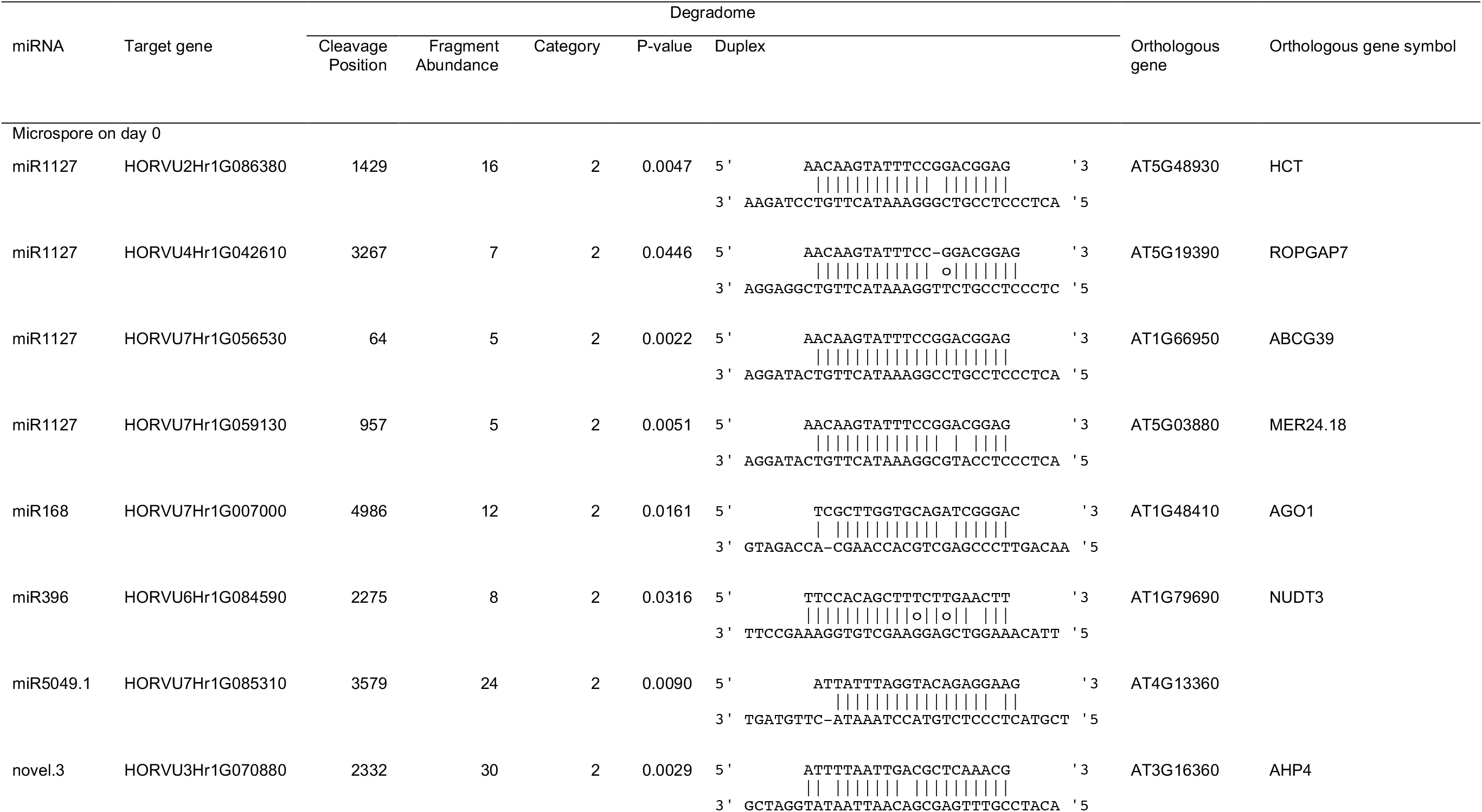
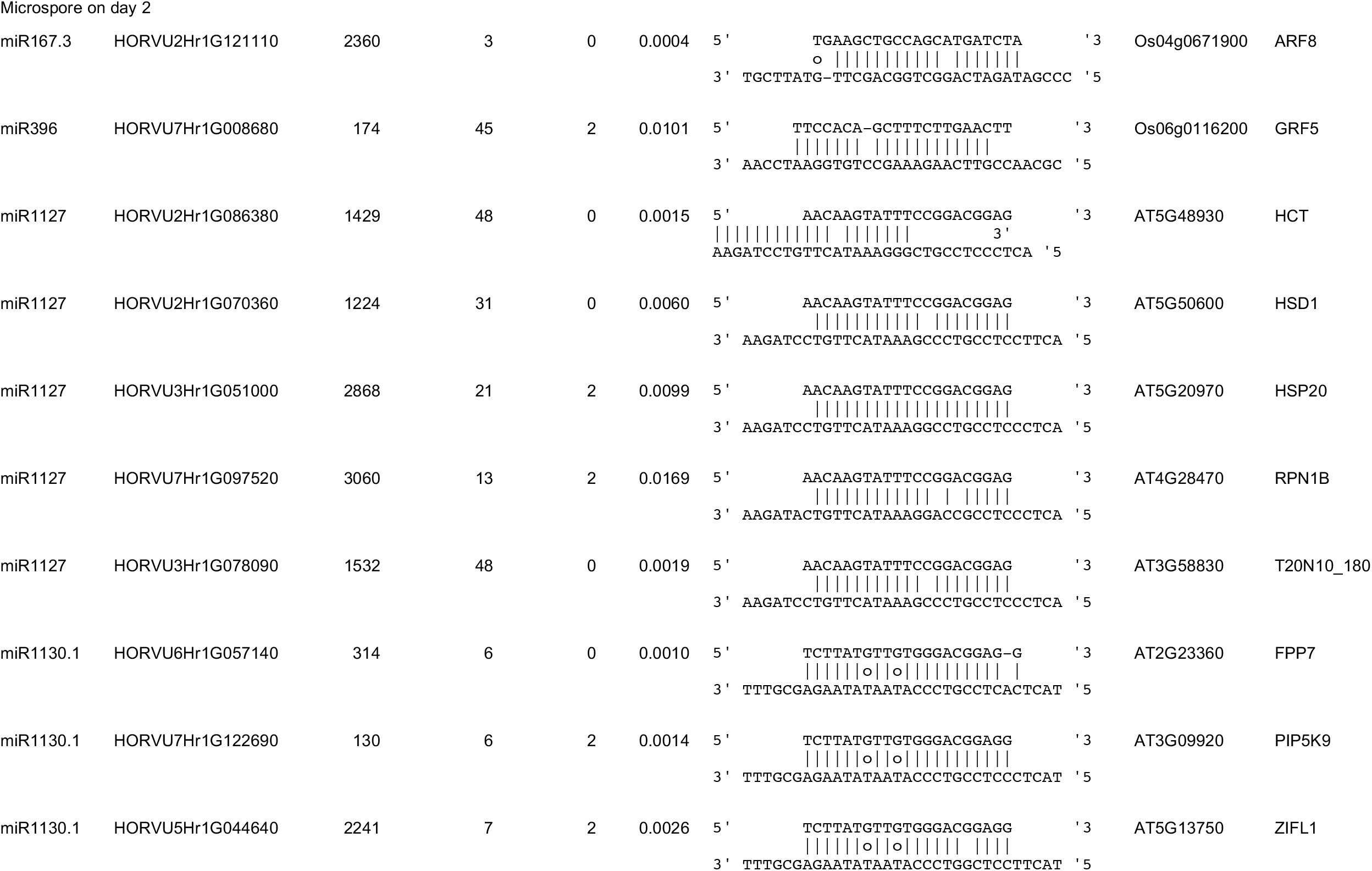
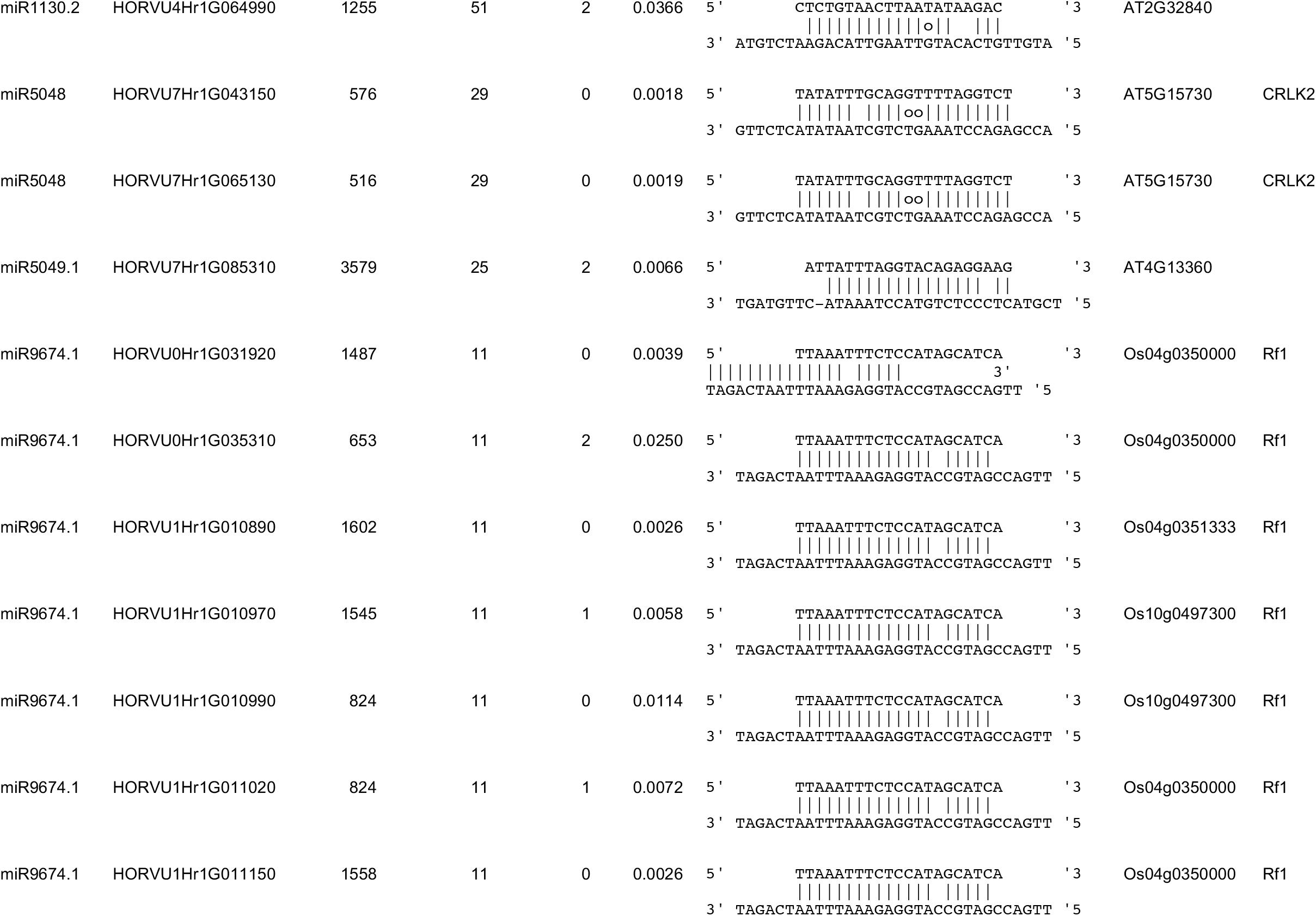
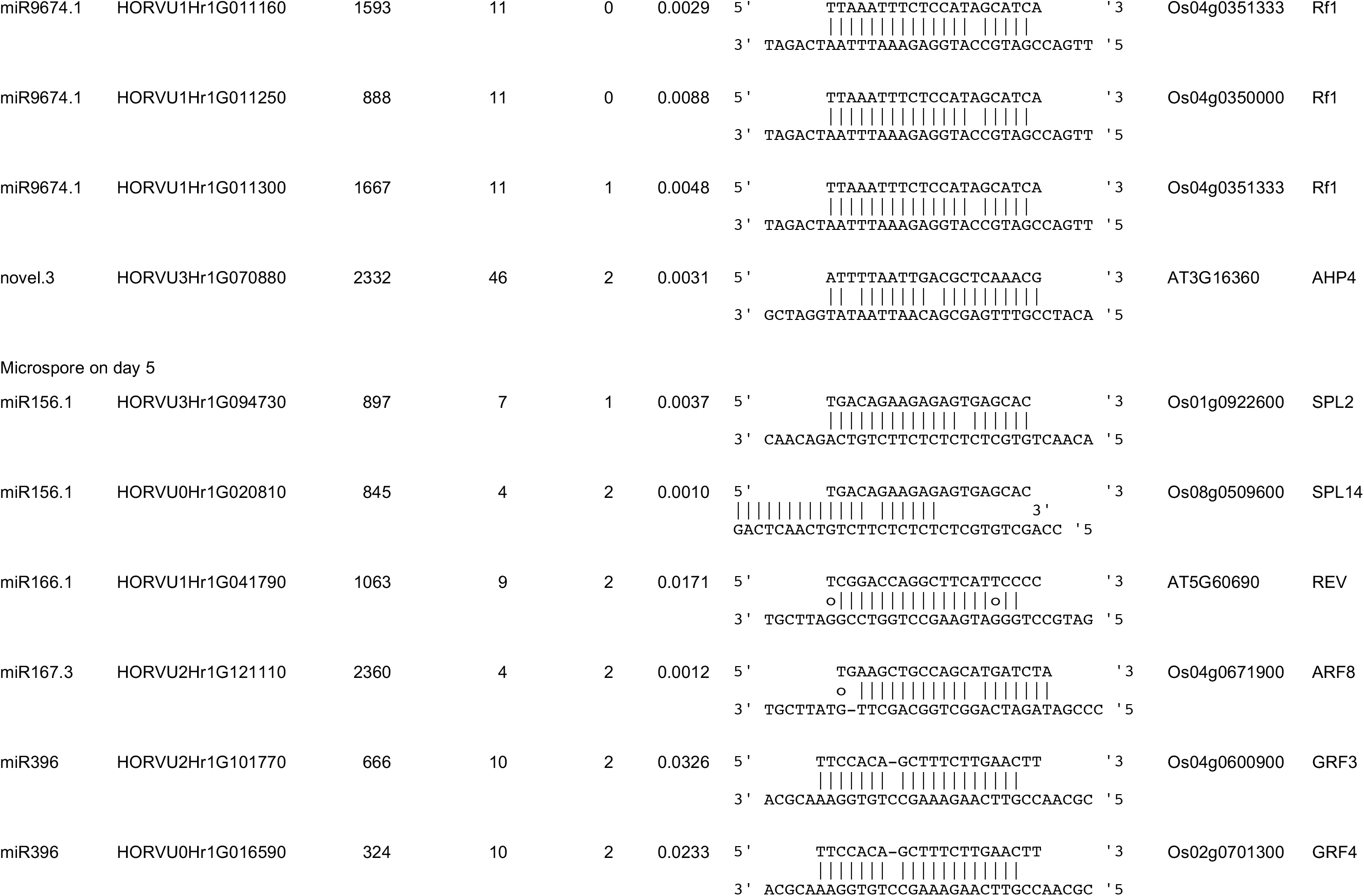
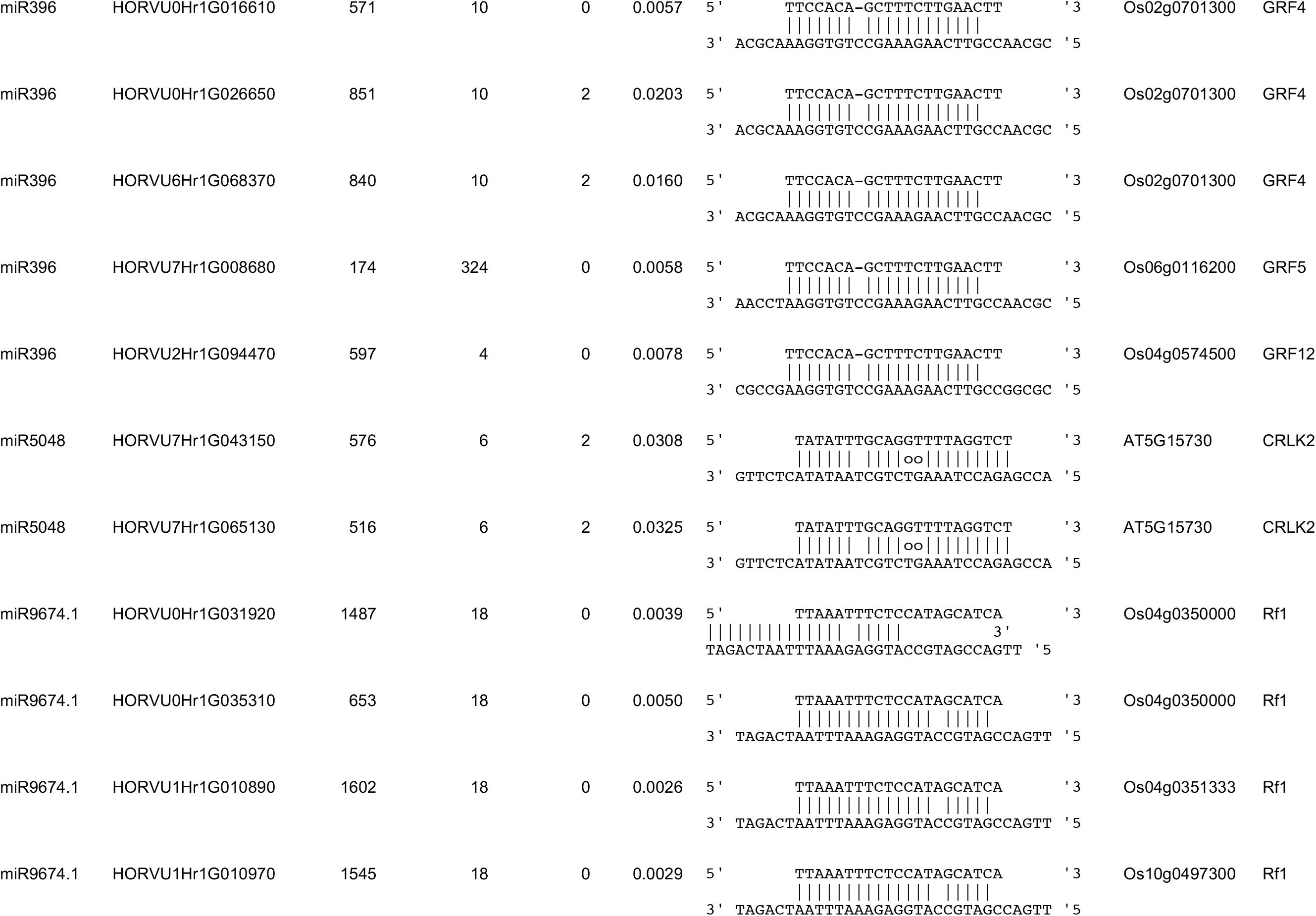
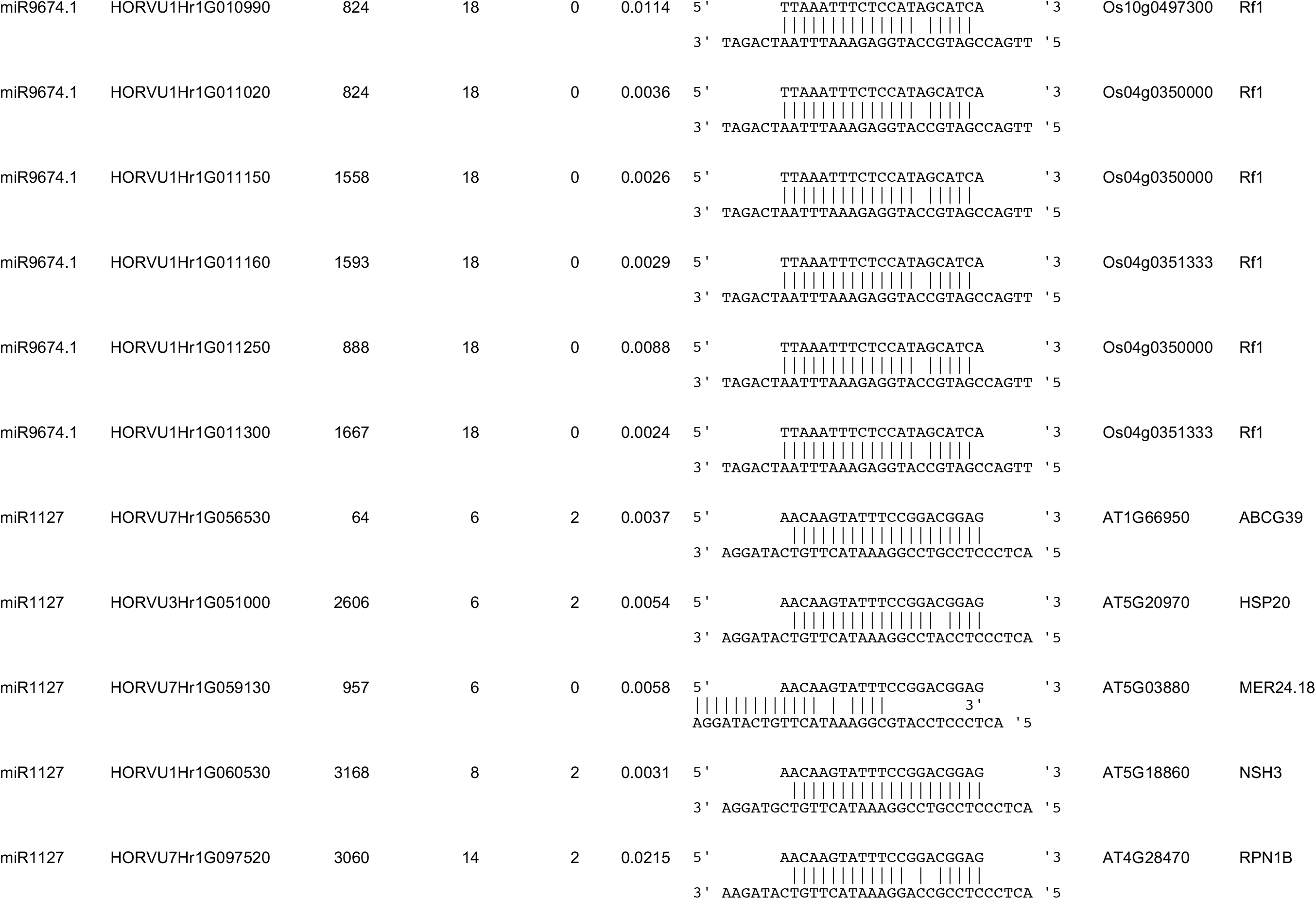
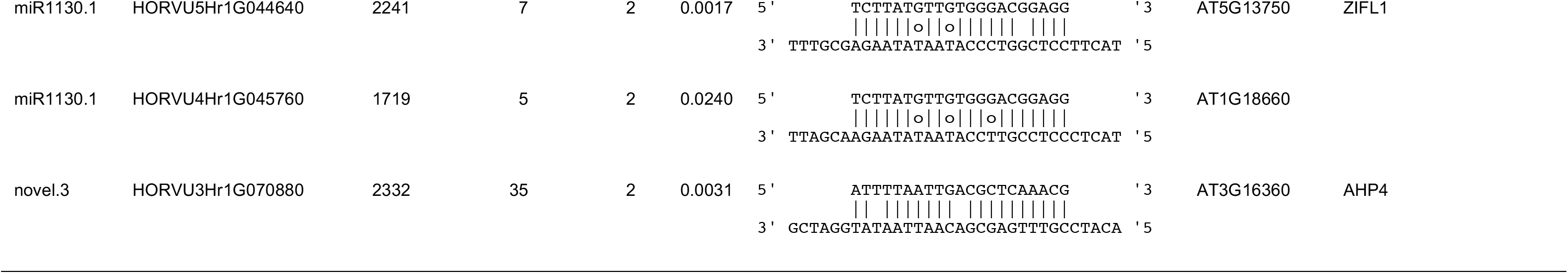
Detail of miRNA targets validated by PARE tags in barley microspores (cv. Gobernadora) undergoing gametic embyrogenesis.

## Notes

### Competing Interest Statement

The authors have declared no competing interest.

